# The RAS GTPase RIT1 compromises mitotic fidelity through spindle assembly checkpoint suppression

**DOI:** 10.1101/2020.11.17.386177

**Authors:** Antonio Cuevas-Navarro, Richard Van, Alice Cheng, Anatoly Urisman, Pau Castel, Frank McCormick

## Abstract

The spindle assembly checkpoint (SAC) is an evolutionarily conserved safety mechanism that maintains genomic stability. However, despite the understanding of the fundamental mechanisms that control the SAC, it remains unknown how signaling pathways directly interact with and regulate the mitotic checkpoint activity. In response to extracellular stimuli, a diverse network of signaling pathways involved in cell growth, survival, and differentiation are activated and this process is prominently regulated by the Ras family of GTPases. Here we show that RIT1, a Ras-related GTPase, is essential for timely progression through mitosis and proper chromosome segregation. Furthermore, pathogenic levels of RIT1 silence the SAC, accelerate transit through mitosis, and promote chromosome segregation errors through direct association with SAC proteins MAD2 and p31^comet^. Our results highlight a unique function of RIT1 compared to other Ras GTPases and elucidate a direct link between a signaling pathway and the SAC through a novel regulatory mechanism.

**Graphical Abstract:** 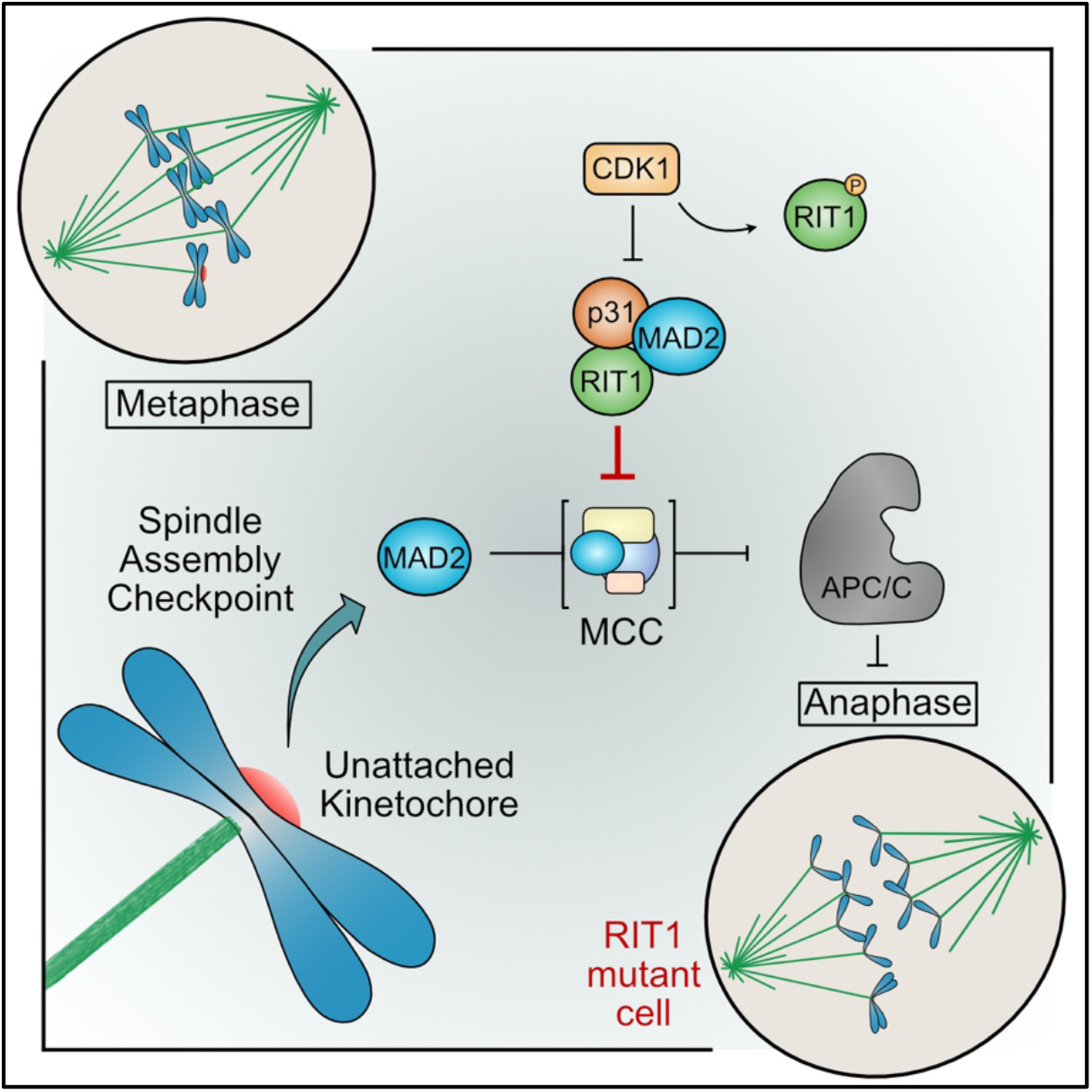

## INTRODUCTION

The spindle assembly checkpoint (SAC) functions as a sensor of unattached kinetochores that delays mitotic progression into anaphase until proper chromosose segregation is guaranteed. Disruptions to this safety mechanism lead to genomic instability and aneuploidy, which serve as the genetic cause of embryonic demise, congenital birth defects, intellectual disability, and cancer (Ben-David and Amon, 2020; Oromendia and Amon, 2014). RIT1 is a Ras-related small guanosine triphosphatase (GTPase) that regulates cell survival and stress response (Van et al., 2020) and mutations in its gene have been identified as oncogenic drivers of lung adenocarcinoma and etiologic factors of Noonan syndrome (Aoki et al., 2013; Berger et al., 2014; Yaoita et al., 2016). A distinctive feature of RIT1, and its paralog RIT2, is the lack of C-terminal prenylation (Wes et al., 1996). Despite this, the RIT1 C-terminal tail mediates plasma membrane association through electrostatic interactions with phospholipids (Heo et al., 2006). RIT1 has a unique set of effector proteins but shares activation of the MAPK pathway with other Ras GTPases (Rodriguez-Viciana et al., 2004; Simanshu et al., 2017; Van et al., 2020). However, due to the lack of identified cognate GTPase activating protein (GAP) or exchange factor (GEF) enzymes, regulation of the RIT1 GTPase cycle remains unclear (Van et al., 2020). Nonetheless, RIT1 abundance and activity is regulated at the protein level through proteasomal degradation, a mechanism that is mediated by the adaptor protein LZTR1 and the E3 ubiquitin ligase Cullin 3 (Castel et al., 2019). While RIT1’s role in Noonan syndrome is likely mediated by the dysregulation of the MAPK pathway, a pathognomonic sign of the disorder, its role in normal cells and in malignancies is less clear. Here, we describe an unprecedended role for a Ras GTPase; the direct interaction of RIT1 with the SAC to promote its inhibition.

## RESULTS

### RIT1 forms a complex with SAC proteins MAD2 and p31^comet^

To characterize the RIT1 interactome, we performed an affinity purification-mass spectrometry screening in mammalian cells (**Figure 1A**). Previously identified interactors included LZTR1, and calmodulin-related proteins CALM2 and CALML3 (Castel et al., 2019; Wes et al., 1996). Our screen also identified two novel binding partners: MAD2 (MAD2L1) and p31^comet^ (also known as MAD2L1-binding protein). RIT1 pull-down assays not only validated the interaction but also revealed that MAD2 and p31^comet^ are unique binding partners of RIT1 that do not interact with other Ras GTPases (**Figures S1A and S1B**).

**Figure 1.**
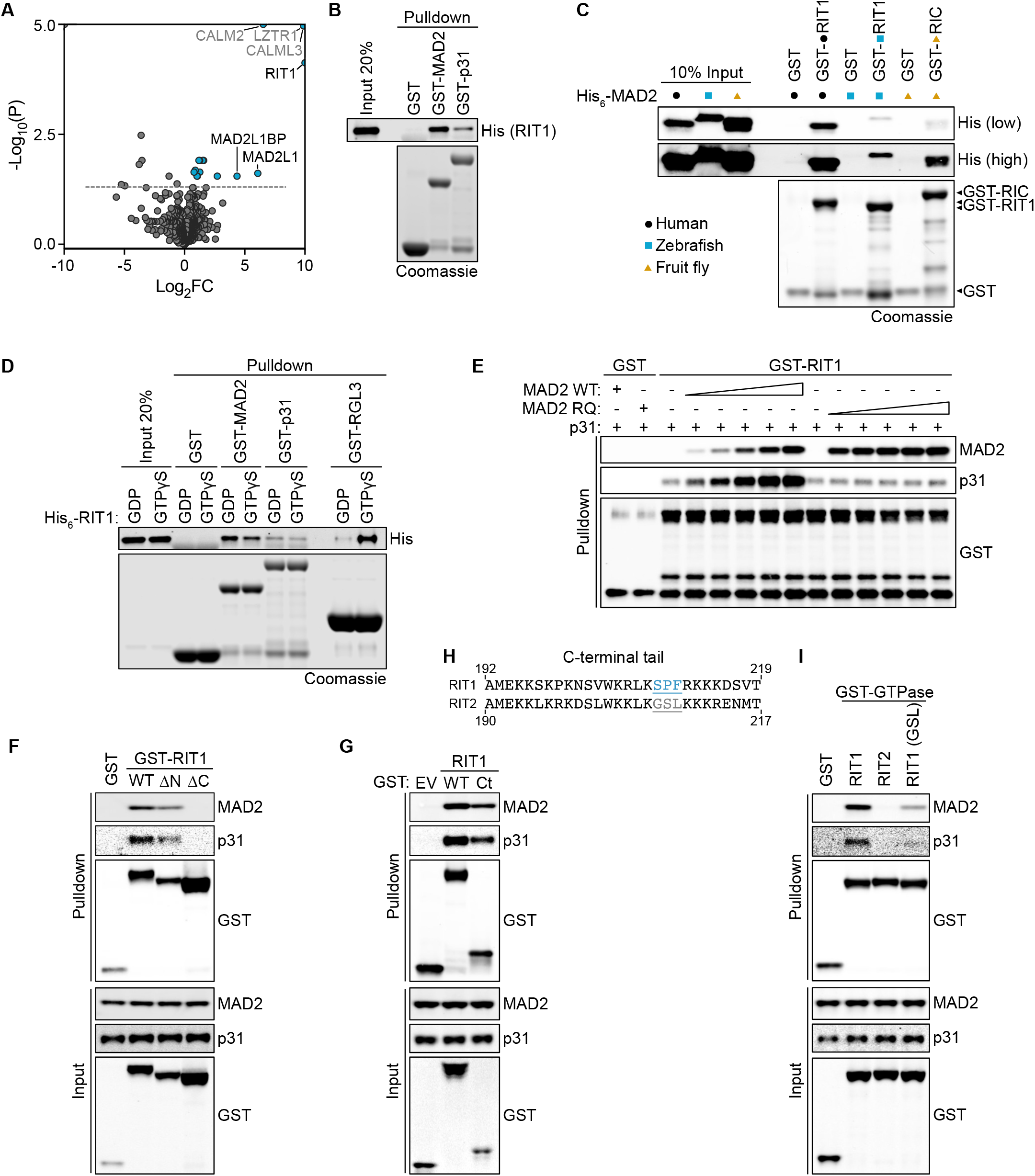
RIT1 interacts directly with the SAC proteins MAD2 and p31^comet^. (**A**) Proteins obtained from lysates of HEK-293T cells transfected with FLAG-RIT1 or FLAG empty vector control were immunoprecipitated and analysed by LC-MS/MS. Volcano plot shows enrichment of proteins detected in FLAG-RIT1 precipitates across three biologically independent repeats. MAD2L1 and MAD2L1BP correspond to the gene names of MAD2 and p31^comet^, respectively. Log_2_ fold change (Log_2_FC) and −Log_10_ adjusted p-value (−Log_10_(P)) were capped at 10 and 5.0, respectively. Dashed line represents p-value of 0.05. (**B**) GST pulldown assay with indicated recombinant purified GST-tagged proteins and His_6_-RIT1. (**C**) GST pulldown assay with 0.1 μM indicated recombinant purified GST-tagged RIT1 or RIC proteins and 0.1 μM MAD2 orthologs. (**D**) GST pulldown assay as in (B), with His_6_-RIT1 protein loaded with GDP or GTPγS. GST-RGL3 serves as a positive control for nucleotide loading due to the GTP-dependent nature of the RIT1-RGL3 interaction. (**E**) GST pulldown assay with 0.1 μM recombinant GST or GST-RIT1 incubated with 0.5 μM p31^comet^ and titration of MAD2 WT or the dimerization and p31^comet^ binding deficient mutant MAD2 R133E/Q134A (RQ) (0, 0.1, 0.2, 0.4, 0.8 or 1.6 μM). (**F**, **G, I**) Proteins precipitated from extracts of HEK-293T cells transfected with GST, GST-RIT1, or GST-RIT1 mutant constructs. Immunoblots were probed for endogenous MAD2 and p31^comet^. EV, empty vector. Ct, C-terminal (192-219). (**H**) Alignment of RIT1 and RIT2 C-terminal extensions. (**I**) RIT1 (GSL), RIT1 construct with residues 209-211 replaced with corresponding RIT2 residues 207-209 (GSL). See also Figure S1.

MAD2 plays a critical role in SAC signal amplification through its association with its ligand MAD1 at unattached kinetochores, a step that catalyzes the formation of the SAC effector, the mitotic checkpoint complex (MCC) (Musacchio, 2015). The MCC, which includes MAD2 bound to its weaker affinity ligand CDC20, inhibits anaphase entry until the SAC is satisfied. In contrast, p31^comet^ inhibits the formation of MCC by binding MAD1-bound MAD2 at unattached kinetochores (Hagan et al., 2011; Xia et al., 2004). Additionally, p31^comet^ participates in the removal of MAD2 from MCC by serving as an adaptor for the AAA ATPase TRIP13, which extracts MAD2 and promotes MCC disassembly in an ATP-dependent manner (Westhorpe et al., 2011; Eytan et al., 2014; Wang et al., 2014). MAD2 and p31^comet^ dimerization in cells prompted us to assess whether RIT1 associates with MAD2 and p31^comet^ directly (Habu et al., 2002). Pulldown analysis using recombinant proteins revealed that RIT1 binding to MAD2 and to p31^comet^ is direct and does not depend on MAD2 and p31^comet^ dimerization (**Figure 1B**). Furthermore, the RIT1-MAD2 interaction is conserved among vertebrate and invertebrate orthologs (**Figure 1C**). To determine whether binding to MAD2 or p31^comet^ is regulated by RIT1’s GTPase cycle, we performed a direct binding assay using recombinant human RIT1 loaded with GDP or GTPγS, a non-hydrolyzable GTP analogue. This revealed that both interactions are independent on the guanosine nucleotide-loaded state of RIT1 (**Figure 1D**). Consistent with these observations, binding to MAD2 and p31^comet^ is not influenced by disease-associated RIT1 mutations (**Figure S1C**). These results suggested that the binding interface lies outside of the switch I and switch II domains of RIT1 that are sensitive to GDP/GTP binding, and, hence, MAD2 and p31^comet^ are not typical RIT1 effector proteins.

The structural similarity between MAD2 and p31^comet^ and our evidence for direct binding to RIT1 predicted either a competitive binding model, in which a single molecule of RIT1 interacts with either MAD2 or p31^comet^, or a non-competitive binding model in which RIT1 binds to MAD2 and p31^comet^ simultaneously (Mapelli et al., 2007; Yang et al., 2007). To discern between these two models, we used a competitive binding assay in which titration of MAD2 WT or RQ, a dimerization and p31-binding deficient mutant, failed to suppress RIT1-p31^comet^ binding (De Antoni et al., 2005; Mapelli et al., 2006) (**Figure 1E**). These data support a non-competitive binding model. Notably, titration of MAD2 exerts a cooperativity effect on RIT1-p31^comet^ binding that is dependent on MAD2 and p31^comet^ dimerization, a behavior that was further observed under analytical size exclusion chromatography (SEC) (**Figure S1D-S1I**). Consistent with previous reports, we observed spontaneous dimerization of bacterially expressed MAD2 (Fang et al., 1998; Sironi et al., 2001) (**Figure S1D**). Incubation of full-length RIT1, which elutes as a monomer, produced a shift in elution volume consistent with an association between RIT1 and dimeric MAD2 (**Figures S1E and S1F**). The weaker affinity interaction between RIT1 and p31^comet^ (**Figure 1B)** was not detected by SEC (**Figures S1G and S1H**). However, incubation of RIT1, MAD2, and p31^comet^ produced a high molecular-weight peak containing all three proteins, suggesting that RIT1, MAD2 and p31^comet^ assemble into a multimeric complex *in vitro* and that RIT1-p31^comet^ binding may be stabilized by MAD2 (**Figure S1I**).

Due to the high degree of similarity between the RIT1 G-domain and that of other Ras GTPases, particularly its paralog RIT2, we hypothesized that MAD2 and p31^comet^ interact mainly through the N-terminal or C-terminal extensions of RIT1 (Lee et al., 1996). We analyzed RIT1 deletion mutants of these extensions and demonstrated that the C-terminal domain is necessary and sufficient for MAD2 and p31^comet^ binding (**Figures 1F, 1G, and S1J**). Consecutive C-terminal truncations allowed us to identify residues 209-211 as critical for MAD2 and p31^comet^ binding (**Figures S1K and S1L**) and, when mutated to corresponding RIT2 residues 207-209 (GSL), the interaction was abrogated (**Figures 1H and 1I**).

### RIT1 interaction with MAD2 and p31^comet^ is regulated by CDK1 phosphorylation

In addition to mediating its interaction with MAD2 and p31^comet^, the RIT1 C-terminus is responsible for its association with the plasma membrane (PM) (Heo et al., 2006). However, because localization of RIT1 at the PM has only been reported in interphase cells, we sought to determine RIT1 subcellular localization during mitosis. We observed difuse cytoplasmic distribution of RIT1 as cells enter mitosis and as they progress through metaphase, after which RIT1 rapidly translocates to the PM (**Figures 2A and 2B**). Correspondingly, a predominant cytoplasmic distribution of endogenous RIT1, but not HRAS, was detected in mitotic cell lysates (**Figure 2C**). Our data indicates that diffusion of RIT1 between the PM and cytoplasm occurs during cell cycle and can therefore influence its association with MAD2 and p31^comet^.

**Figure 2.**
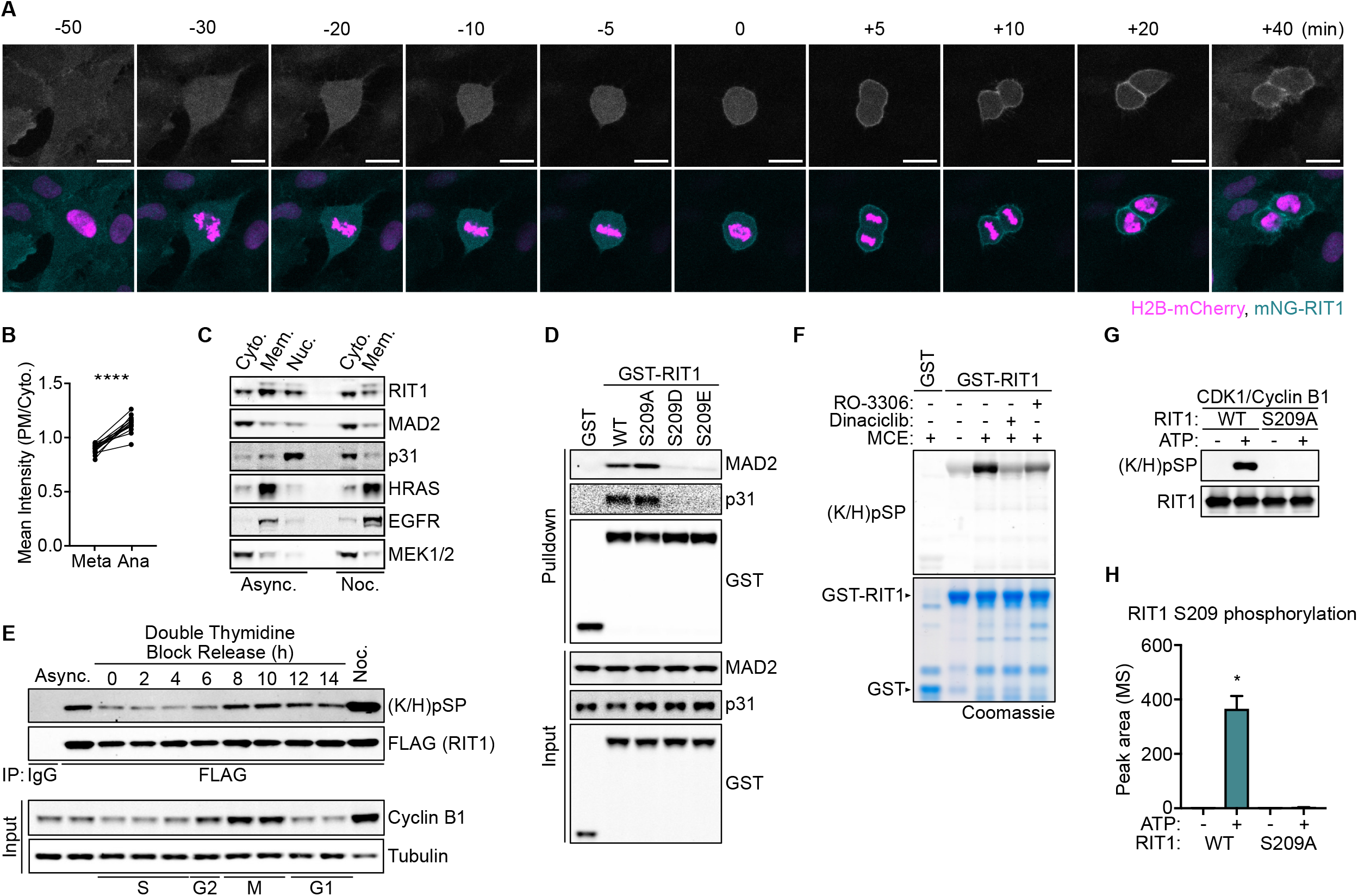
RIT1 interaction with MAD2 and p31^comet^ is regulated by CDK1 phosphorylation. (**A**) hTERT-RPE1 cell stably expressing mNeonGreen (mNG)-RIT1 and Histone H2B-mCherry undergoing mitosis imaged at 5 min intervals. Anaphase onset set to t = 0 min. Scale bar, 20 μm (**B**) Quantification of the ratios of plasma membrane (PM) and cytoplasmic (Cyto.) mNG-RIT1 signals of cells in (A) during metaphase (Meta, −5 min) and anaphase (Ana, +5 min). Two-sided Student’s paired *t*-test, n = 15, *\*\*\*\*P*≤ 0.0001). (**C**) Immunoblots of subcellular protein fractionation of HeLa cell lysates. Async., asynchronous growing cells. Noc., cells released from G1/S arrest for 4 hours then treated with 100 ng/ml nocodazole for 10 hours. (**D**) Protein precipitated from extracts of HEK-293T cells transfected with GST or GST-RIT1 constructs. Immunoblots were probed for endogenous MAD2 and p31^comet^. (**E**) HeLa cells stably expressing FLAG-RIT1 released from a G1/S phase arrest and lysed at indicated time points. Immunoprecipitated proteins were probed for RIT1 S209 phosphorylation by immunoblotting. Async., asynchronous growing cells. Noc., cells released from G1/S arrest for 4 hours then treated with 100 ng/ml nocodazole for 10 hours. (**F**) Detection of RIT1 S209 phosphorylation on bacterially expressed GST-RIT1 protein incubated with mitotic cell extract (MCE) treated with 1 μM Dinaciclib, 10 μM RO-3306, or DMSO control. (**G**) Immunoblot of RIT1 S209 phosphorylation on bacterially expressed RIT1 proteins subjected to an *in vitro* kinase assay with recombinant active CDK1/Cyclin B1. (**H**) MS quantification of phospho-S209 peptides in RIT1 protein incubated with CDK1/Cyclin B1 as in (G). (*n*= 2), Two-sided Student’s *t*-test, data shown as mean, error bars indicate s.e.m, *\*P*≤ 0.05. See also Figure S2.

Because our mass spectrometry revealed that RIT1 is phosphorylated within the C-terminal domain at S209 (**Figure S2A**), we hypothesized that a dynamic regulatory mechanism could control the interaction between RIT1 and MAD2 or p31^comet^. Using phospho-deficient (S209A) and phospho-mimetic (S209D and S209E) mutations we demonstrated that phosphorylation at S209 disrupts RIT1-MAD2/p31^comet^ binding (**Figure 2D**). Additionally, an antibody that detects RIT1 S209 phosphorylation (**Figure S2B**), showed that RIT1 phosphorylation is most abundant during mitosis (**Figure 2E**). To identify the kinase responsible for such phosphorylation, we tested a panel of inhibitors against proline-directed serine/threonine kinases (Hall and Vulliet, 1991) (**Figure S2C**). Inhibition of cyclin-dependent kinase (CDK) activity in prometaphase-arrested cells led to a reduction of RIT1 S209 phosphorylation. Furthermore, in a cell-free assay, using mitotic cell extracts, CDK1 inhibition significantly reduced phosphorylation of recombinant RIT1 (**Figure 2F**). To determine whether RIT1 is a direct substrate of CDK1/Cyclin B1, we performed *in vitro* kinase assays using recombinant purified proteins. Phosphorylation of RIT1 S209 by CDK1/Cyclin B1 was detected by immunoblotting and confirmed by mass spectrometry (**Figures 2G and 2H).**These findings suggest Cyclin B1/CDK1 phosphorylate RIT1 during mitosis, which coincides with the cell cycle pattern of Cyclin B1 expression and CDK1 activity (Lindqvist et al., 2009). We propose that CDK1 phosphorylation regulates the association of RIT1 with MAD2 and p31^comet^ in a cell cycle-dependent manner.

### RIT1 inhibits the SAC and promotes chromosome segregation errors

SAC signaling is tightly regulated and amplified by the catalysed conversion of MAD2 from its open state (O-MAD2) to its closed conformational state (C-MAD2), which associates preferentially with CDC20 and the MCC (Chao et al., 2012; De Antoni et al., 2005; Luo et al., 2000; Musacchio and Salmon, 2007). Amplification of the SAC is silenced by direct binding of p31^comet^ to MAD2 (Westhorpe et al., 2011; Yang et al., 2007). As such, MAD2 and p31^comet^ control the duration of the SAC and, in turn, the duration of mitosis. This prompted us to examine whether RIT1, through its direct association with MAD2 and p31^comet^, can influence the SAC. Depletion of RIT1 by RNAi or by CRISPR-mediated knockout resulted in prolonged mitotic progression (**Figures 3A, 3B, S3A-S3D).**Moreover, pharmacological inhibition of the SAC rescued the effect of RIT1 depletion, indicating that RIT1 affects mitosis upstream of the SAC (Santaguida et al., 2010). Furthermore, loss of RIT1 increased the rate of chromosome segregation errors (**Figure 3C**), suggesting that RIT1 is not only essential for timely progression through mitosis, but that dysregulation of RIT1 protein levels disrupt proper SAC function.

**Figure 3.**
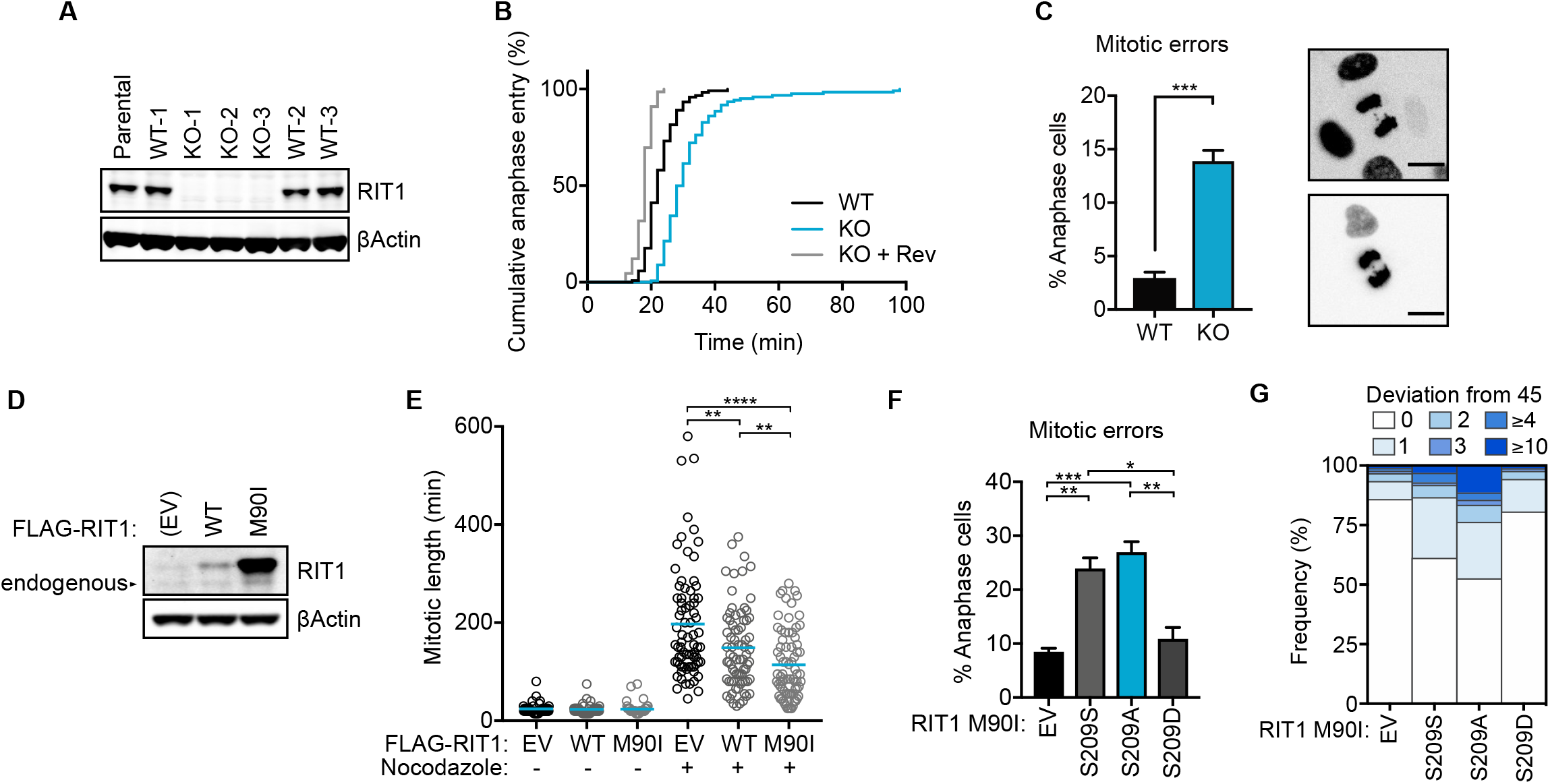
RIT1 inhibits the SAC and promotes chromosome segregation errors. (**A**) Immunoblots of lysates from WT and RIT1 KO hTERT-RPE1 clones. (**B**) Comparison of mitotic transit times between WT and RIT1 KO hTERT-RPE1 cells assessed by timelapse microscopy. Time measured from nuclear envelope breakdown (NEBD). Rev, cells treated with 1 μM Reversine. WT (*n*=119), KO (*n*= 122), KO + Rev (*n*= 66). (**C**) Quantification of mitotic errors in hTERT RPE1. Data represent three independent replicates with three WT clones and three KO clones. Error bars indicate s.e.m. (**D**) Immunoblots of lysates from U2-OS stably expressing indicated constructs. (**E**) Duration of mitotic length (NEBD - anaphase onset) assessed by time-lapse microscopy in U2-OS cells stably expressing indicated proteins treated with 15 ng/ml nocodazole (*n*= 75) or DMSO control (*n*= 50). Two-sided Student’s *t*-test, error bars indicate s.e.m., \**P*≤ 0.05, \*\**P*≤ 0.01, \*\*\**P*≤ 0.001, \*\*\*\**P*≤ 0.0001. (**F**) Comparison of chromosome segregation error rates in HCT-116 cells stably expressing indicated constructs. EV, empty vector. Data represent three biologically independent repeats. Two-sided Student’s *t*-test, error bars indicate s.e.m., \**P*≤ 0.05, \*\**P*≤ 0.01, \*\*\**P*≤ 0.001, \*\*\*\**P*≤ 0.0001. (**G**) Metaphase spread assay compares frequency of aneuploidy, determined by a chromosome count other than the modal number, 45, in HCT-116 cells stably expressing EV (*n*= 92), S209S (*n*= 98), S209A (*n*= 97), or S209D (*n*= 88). For (B, E, G), *n* indicates the number of cells or metaphase spreads counted and data shown is representative of at least two biologically independent experiments. See also Figure S3.

To further assess the effect of RIT1 on the SAC, we expressed RIT1 M90I, a pathogenic variant that is insensitive to protein degradation, resulting in increased expression levels (**Figure 3D**) (Castel et al., 2019). Ectopic expression of RIT1 WT or M90I partially overrides the pharmacologically-induced SAC response in asynchronously growing cells (**Figures 3E and S3E**). To determine whether the RIT1-mediated effect on the SAC is dependent on its association with MAD2 and p31^comet^, we expressed the RIT1 phosphomimetic mutants unable to bind these proteins in cells (**Figures S3F and S3G**). Abolishing MAD2 and p31^comet^ binding rescued suppression of the SAC mediated by RIT1 M90I. We did not observe a discernible difference in basal MAPK activation between these mutants, ruling out the possibility that the rescue effect exhibited by the phospho-mimetic RIT1 mutants was due to altered MAPK signaling (**Figure S3H**).

A weakened SAC may allow precocious anaphase entry that results in chromosome instability and missegregation (Dobles et al., 2000; Hanks et al., 2004; Iwanaga et al., 2007; Li et al., 2009; Michel et al., 2001). Therefore, we examined whether RIT1-mediated suppression of the SAC promotes chromosome segregation errors in HCT-116, a cancer cell line with near diploid karyotype that exhibits low chromosomal instability (Lengauer et al., 1997). Ectopic expression of RIT1 M90I significantly increased the rate of mitotic errors, including lagging and bridging chromosomes, in a MAD2- and p31^comet^-binding dependent manner (**Figures 3F and S3I).**Consequently, we observed an increased rate of aneuploidy in cells expressing RIT1 M90I, but not in cells expressing the mutant that is unable to bind MAD2/p31^comet^ (**Figures 3G and S3J)**. These results demonstrate that increased levels of RIT1 led to compromised mitotic fidelity as a result of direct interaction with MAD2 and p31^comet^.

### RIT1 inhibits MAD2 association with SAC complexes and promotes degradation of APC/C substrates

Proper SAC signaling relies on the recruitment of cytosolic O-MAD2 to unattached kinetochores via a high-affinity interaction with MAD1 that drives the conversion of O-MAD2 to C-MAD2 (De Antoni et al., 2005; Mapelli et al., 2007; Sironi et al., 2002). MAD1 binding mediates the dimerization of MAD1-bound C-MAD2 with a second molecule of MAD2. The latter step stabilizes the second molecule of MAD2 in an intermediate state that promotes its association with CDC20 (Hara et al., 2015; Luo et al., 2002). To directly test whether RIT1 inhibits the association of MAD2 with CDC20 and MAD1, we performed competitive pulldown assays to test mutual exclusivity between RIT1-MAD2 and MAD2-CDC20/MAD1 binding (**Figures 4A and 4B**). MAD2 binding peptide 1 (MBP1), a high-affinity synthetic peptide that mimics the binding of MAD2 to the MAD2 interaction motifs (MIM) of CDC20 and MAD1, abolished MAD2-RIT1 binding (Luo et al., 2002). Conversely, titration of full-length RIT1 protein reduced binding of MAD2 to CDC20 MIM (residues 111-138) beads (Mondal et al., 2006). These results suggest that RIT1 competes with CDC20 and MAD1 for MAD2 binding. However, since binding to MBP1 or CDC20^111-138^ drives the conversion of O-MAD2 to C-MAD2, an alternative explanation may be that RIT1 preferentially binds to O-MAD2 (Luo et al., 2002). To distinguish between these two models, we assessed binding of RIT1 with MAD2 mutants that stabilize MAD2 in either its open or closed conformational state (Mapelli et al., 2007) (**Figure 4C**). C-MAD2 stabilizing mutants retained their interaction to RIT1, whereas O-MAD2 stabilizing mutants failed to bind RIT1, with the exception of MAD2 LL, which can adopt a closed conformation state in the presence of MIM ligand (Hara et al., 2015). Furthermore, expression of a MAD2 phospho-mimetic mutant in cells that adopts the O-MAD2 conformer fails to bind RIT1 in pulldown assays (Kim et al., 2010) (**Figure S4A**). These results demonstrate that RIT1 exhibits preferential binding to C-MAD2 over O-MAD2 and that RIT1 competes with CDC20 and MAD1 for MAD2 binding. We posit that RIT1 and CDC20/MAD1 compete for the same interface on MAD2, which would predict that RIT1 binding may also drive the conversion of O-MAD2 to C-MAD2. To test this hypothesis, we incubated MAD2 protein with excess RIT1 peptide corresponding to residues 202-216, then separated by gel filtration (**Figure 4D**). In the presence of excess RIT1 peptide, MAD2 protein fails to dimerize, similar to previous reports in which the use of excess CDC20 peptide disrupted MAD2 dimerization by saturating all molecules into their closed conformational state (DeAntoni et al., 2005; Sironi et al., 2001). In contrast, incubation of MAD2 with S209 phosphorylated RIT1 peptide did not disrupt the formation of MAD2 dimers (**Figure 2D**). RIT1 preferential binding to C-MAD2 and dimerization of p31^comet^ with C-MAD2 suggests that their oligomerization produces a RIT1-C-MAD2-p31^comet^ complex and, hence, would explain the increased RIT1-p31^comet^ affinity observed in the presence of MAD2 (**Figure 1E**). To investigate the effect of RIT1 on MAD2 and CDC20 binding in the presence of p31^comet^, we conducted competitive pulldown assays. Based on the cooperativity effect exhibited by the RIT1-MAD2-p31^comet^ complex, the addition of p31^comet^ cooperated with RIT1-mediated inhibition of MAD2 and CDC20 binding (**Figures S4B and S4C**). Furthermore, RIT1 and p31^comet^ cooperation was dependent on MAD2-p31^comet^ binding. These results suggest that RIT1 may cooperate with p31^comet^ to extract MAD2 from the MCC and promote its disassembly.

**Figure 4.**
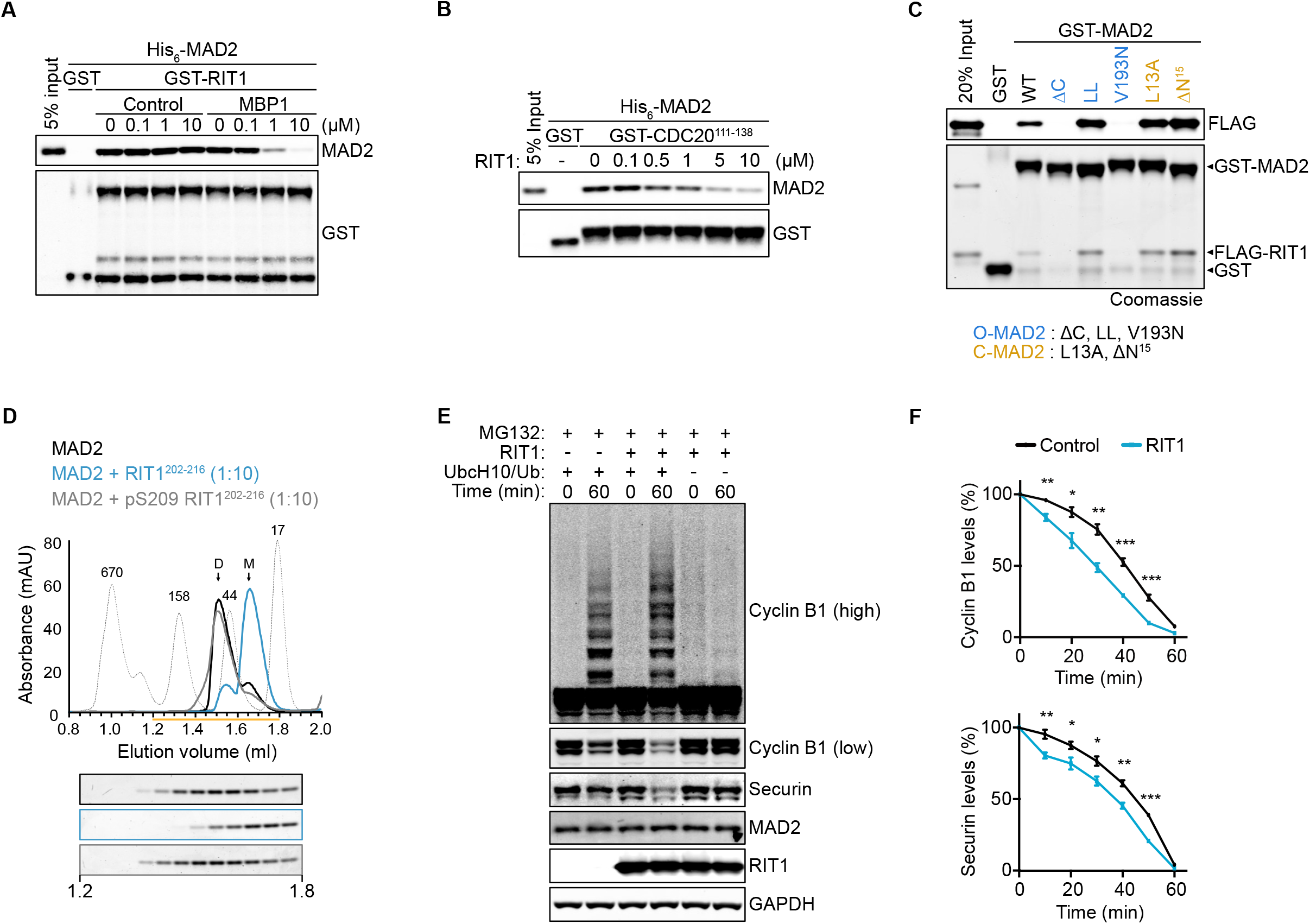
RIT1 inhibits MAD2 association with SAC complexes and promotes degradation of APC/C substrates. (**A**, **B**) Immunoblots of precipitated proteins from an equilibrium competition pulldown assays with 0.2 μM recombinant (A) GST-RIT1 or (B) GST-CDC20 111-138 protein incubated with 0.2 μM His_6_-MAD2 and titrating amounts of (A) MAD2 binding peptide 1 (MBP1) or a control peptide or (B) full-length RIT1 protein. (**C**) Pulldown assay with 0.5 μM recombinant GST or GST-MAD2 proteins incubated with 0.5 μM FLAG-RIT1 protein. Precipitated proteins were separated by SDS-PAGE for immunoblot or Coomassie staining. (**D**) Elution profile of MAD2 protein incubated with or without indicated peptides at 1:10 molar ratio for 1 hour at 25°C prior to gel filtration. The contents of 12 consecutive 50 μl fractions eluting between 1.2 ml to 1.8 ml are shown. (**E**) Immunoblots of samples from APC/C ubiquitination assay with and without 10 μM RIT1. To prevent degradation of ubiquitinated proteins, mitotic cell extracts were incubated with 10 μM MG132 for 1 hour on ice, prior to supplementation with additional ubiquitination assay components. (**F**) Comparison of Cyclin B1 (top panel) and Securin (bottom panel) degradation in APC/C activity assays supplemented with or without recombinant RIT1 protein (10 μM). Data represent four biologically independent repeats. Two-sided Student’s *t*-test, data points are mean ± s.e.m., \**P*≤ 0.05, \*\**P*≤ 0.01, \*\*\**P*≤ 0.001. See also Figure S4.

Inhibition of mitotic progression by the SAC is achieved through suppression of the multisubunit ubiquitin ligase anaphase-promoting complex/cyclosome (APC/C) by the MCC (Primorac and Musacchio, 2013). We reasoned that RIT1 inhibition of MAD2-CDC20 association may promote APC/C activity. Therefore, we measured *in vitro* ubiquitination and degradation of the main APC/C substrates, Cyclin B1 and Securin, in MCC active mitotic cell extracts isolated from RIT1 knockout cells (Chang and Barford, 2014). Adding recombinant RIT1 protein to MCC active mitotic cell extracts increased ubiquitination and degradation of Cyclin B1 and Securin, suggesting increased APC/C activity in the presence of RIT1, likely due to relieved MCC inhibition (**Figures 4E, 4F, S4D, S4E**).

## Discussion

Altogether, our findings demonstrate that RIT1 regulates mitotic fidelity through a direct complex formation with MAD2 and p31^comet^ that results in suppression of the SAC response. This unique function may be attributed, in part, to the distinct regulation of RIT1 localization; the absence of C-terminal prenylation that anchors Ras GTPases to the PM (Simanshu et al., 2017) may promote the diffusion of RIT1 between PM and cytoplasm, a potential requisite for its association with MAD2 and p31^comet^. Characterization of this complex revealed a regulatory mechanism mediated by CDK1 activity during mitosis. CDK1/Cyclin B1 inhibits the formation of RIT1-MAD2-p31^comet^ complexes through RIT1 C-terminal tail phosphorylation. CDK1 orchestrates mitotic progression through phosphorylation of various substrates and is most active during prometaphase (Hochegger et al., 2008). In line with this, we can deduce that CDK1/Cyclin B1 suppresses RIT1-mediated SAC inhibition during the early stages of mitosis when SAC signaling is essential for proper chromosome segragation (Musacchio, 2015). Pathogenic levels of RIT1 suppress SAC signaling and we speculate that overabundant RIT1 escape adequate CDK1 phosphorylation achieved under physiological RIT1 levels, resulting in a weakened SAC response. To fully understand the properties of RIT1 as an oncogenic driver, it will be paramount to investigate whether SAC silencing contributes to the pathogenesis of *de novo* RIT1 mutations that compromise RIT1 degradation, as this may provide an avenue for therapeutic intervention (Castel et al., 2019).

We further demonstrate that MAD2 and p31^comet^ are not typical RIT1 effector proteins and that the RIT1 C-terminal tail is sufficient for binding; however, the role of its GTPase activity on SAC regulation remains to be evaluated. It is tempting to speculate that RIT1 may provide a direct link between SAC regulation and RIT1 effector pathways involved in cell survival and stress response (Van et al., 2020). Moreover, one can postulate that the RIT1-SAC signaling axis may have evolved as a mechanism that modulates SAC activity in response to mitogenic and stress signals. While previous reports have implicated pathogenic Ras GTPase signaling in the dysregulation genomic stability (Hingorani et al., 2005; Kamata and Pritchard, 2011; Yang et al., 2013), our results show a direct link between the SAC and a member of the Ras GTPase family, providing a novel example of the evolutionary adaptation of a signaling molecule for the regulation of a unique but critical cellular pathway.

## SUPPLEMENTARY MATERIALS

Figures S1-S4

## ACKNOWLEDGEMENTS

We thank Dom Esposito and Vanessa Wall for the multisite gateway plasmid toolkit and members of the McCormick lab for their input. Data for this study were acquired at the Nikon Imaging Center at UCSF. This work was supported by a grant from the NCI (R35CA197709 to F.M.). ACN is a fellow of the NSF Graduate Research Fellowship Program (1650113). P.C. work is supported by the NCI (K99CA245122). We thank the UCSF Mass Spectrometry Facility and A. L. Burlingame for providing MS instrumentation support for this project (funded by the NIH grants P41GM103481 and S10OD016229).

## AUTHOR CONTRIBUTIONS

A.C.-N., P.C. and F.M. conceived the project, supervised the research, and wrote the manuscript. A.C.-N. prepared the figures. A.C.-N. and R.V. performed the experiments. A.C. performed chromosome spread analysis. A.C. and A.U. performed MS sample preparation and analysis. All authors commented on and approved the manuscript.

## DECLARATION OF INTERESTS

The authors declare no competing interests.

## STAR★METHODS

### KEY RESOURCES TABLE

| REAGENT or RESOURCE | SOURCE | IDENTIFIER |
| --- | --- | --- |
| Antibodies |  |  |
| Phospho-p44/42 MAPK (Erk1/2) (Thr202/Tyr204) (D13.14.4E) XP Rabbit mAb | Cell Signaling Technology | Cat.# 4370 |
| p44/42 MAPK (Erk1/2) (L34F12) Mouse mAb | Cell Signaling Technology | Cat.# 4696 |
| Phospho-MEK1/2 (Ser217/221) (41G9) Rabbit mAb | Cell Signaling Technology | Cat.# 9154 |
| MEK1/2 (L38C12) Mouse mAb | Cell Signaling Technology | Cat.# 4694 |
| MEK1/2 (D1A5) Rabbit mAb | Cell Signaling Technology | Cat.# 8727 |
| Cyclin B1 (V152) Mouse mAb | Cell Signaling Technology | Cat.# 4135 |
| HA-Tag (C29F4) Rabbit mAb #3724 | Cell Signaling Technology | Cat.# 3724 |
| DYKDDDDK Tag (D6W5B) Rabbit mAb | Cell Signaling Technology | Cat.# 14793 |
| GST (91G1) Rabbit mAb | Cell Signaling Technology | Cat.# 2625 |
| Phospho-CDK Substrate Motif [(K/H)pSP] MultiMab Rabbit mAb | Cell Signaling Technology | Cat.# 9477 |
| PTTG Antibody (DCS-280) Mouse mAb | Santa Cruz Biotechnology | Cat.# sc-56207 |
| GST Antibody (B-14) Mouse mAb | Santa Cruz Biotechnology | Cat.# sc-138 |
| MAD2 Antibody (17D10) Mouse mAb | Santa Cruz Biotechnology | Cat.# sc-47747 |
| RIT1 Rabbit polyclonal | Abcam | Cat.# ab53720 |
| p31comet Antibody, clone E29.19.14 mAb | EMD Millipore | Cat.# MABE451 |
| MAD2 Antibody, Rabbit polyclonal | Bethyl Laboratories | Cat. # A300-301A |
| β-Actin antibody, mAb | Sigma-Aldrich | Cat.# A2228 |
| FLAG M2, mAb | Sigma-Aldrich | Cat.# F1804 |
| Histidine Tagged Antibody, clone HIS.H8, mAb | Sigma-Aldrich | Cat.# 05-949 |
| α-Tubulin antibody, mAb | Sigma-Aldrich | Cat.# T6199 |
| Rabbit IgG (H&L) Antibody DyLight™ 680 Conjugated, Goat polyclonal | Rockland Inc. | Cat.# 611-144-002 |
| Mouse IgG (H&L) Antibody DyLight™ 800 Conjugated Goat Polyclonal | Rockland Inc. | Cat.# 610-145-002 |
| Bacterial and Virus Strains |  |  |
| BL21(DE3) | New England Biolabs | Cat.# C2527H |
| DH5α | McCormick Lab | N/A |
| Stbl3 | McCormick Lab | N/A |
| Chemicals, Peptides, and Recombinant Proteins |  |  |
| Nocodazole | Selleckchem | Cat.# S2775 |
| Reversine | Selleckchem | Cat.# S7588 |
| Dinaciclib | Selleckchem | Cat.# S2768 |
| RO-3306 | Selleckchem | Cat.# S7747 |
| JNK-IN-8 | Selleckchem | Cat.# S4901 |
| CHIR-99021 | Selleckchem | Cat.# S2924 |
| TG003 | Selleckchem | Cat.# S7320 |
| Losmapimod | Selleckchem | Cat.# S7215 |
| Trametinib | Selleckchem | Cat.# S2673 |
| MG132 | Sigma-Aldrich | Cat.# M8699 |
| Thymidine | Sigma-Aldrich | Cat.# T1895 |
| ATP | Sigma-Aldrich | Cat.# A7699 |
| Creatine Phosphate | Sigma-Aldrich | Cat.# 2380 |
| GDP | Sigma-Aldrich | Cat.# G7127 |
| GTPγS | Sigma-Aldrich | Cat.# G8634 |
| MBP1 (SWYSYPPQRAV) | GenScript | N/A |
| Control peptide (SYWPQRAPPSVY) | GenScript | N/A |
| RIT1 (202-216) peptide | GenScript | N/A |
| pS209 RIT1 (202-216) peptide | GenScript | N/A |
| Hexadimethrine bromide | Sigma-Aldrich | Cat.# 107689 |
| Isopropyl β-D-1-thiogalactopyranoside (IPTG) | Sigma-Aldrich | Cat.# I5502 |
| Protease Inhibitor Cocktail | Sigma-Aldrich | Cat.# P8849 |
| TEV-Protease | Sigma-Aldrich | Cat.# T4455 |
| Phosphatase Inhibitor Cocktail 1 | Sigma-Aldrich | Cat.# P2850 |
| Phosphatase Inhibitor Cocktail 2 | Sigma-Aldrich | Cat.# P5726 |
| Protease Inhibitor Cocktail (mammalian cell extracts) | Sigma-Aldrich | Cat.# P8340 |
| ProLong Gold Antifade Mountant with DAPI | Invitrogen | Cat.# P36935 |
| KaryoMAX Colcemid Solution in PBS | Thermo Fisher Scientific | Cat.# 15212012 |
| GlutaMAX | Thermo Fisher Scientific | Cat.# 35050061 |
| active CDK1/Cyclin B1 | SignalChem | Cat.# C22-10G |
| Ubiquitin | Boston Biochem | Cat.# U-100H |
| UbcH10 | Boston Biochem | Cat.# E2-650 |
| Creatine phosphokinase | Sigma-Aldrich | Cat.# C7886 |
| Critical Commercial Assays |  |  |
| MycoAlert PLUS Mycoplasma Detection Kit | Lonza | Cat.# LT07-710 |
| Subcellular Protein Fractionation Kit for Cultured Cells | Thermo Fisher Scientific | Cat.# 78840 |
| NuPAGE 4-12% Bis-Tris Gel | Invitrogen | Cat.# NP0321BOX |
| NuPAGE Sample Loading Buffer | Invitrogen | Cat.# NP0007 |
| Glutathione Sepharose 4B | Cytiva Life Sciences | Cat.# 17075605 |
| ANTI-FLAG M2 Affinity Gel | EMD Millipore | Cat.# A2220 |
| ZipTip with 0.6 μL C18 resin | Millipore | Cat.# ZTC18S008 |
| Gateway LR Clonase II Enzyme Mix | Thermo Fisher Scientific | Cat.# 11791100 |
| Gateway BP Clonase II Enzyme Mix | Thermo Fisher Scientific | Cat.# 11789100 |
| Superdex 200 Increase 3.2/300 column | Cytiva Life Sciences | Cat.# 28990946 |
| RESOURCE Q anion exchange chromatography column | Cytiva Life Sciences | Cat.# 17117701 |
| HisTrap FF (1ml) column | Cytiva Life Sciences | Cat.# 17531901 |
| Superdex 75 Increase 10/300 GL column | Cytiva Life Sciences | Cat.# 29148721 |
| Deposited Data |  |  |
| Experimental Models: Cell Lines |  |  |
| Human: HEK-293T | ATCC | Cat.# CRL-3216 |
| Human: HeLa | ATCC | Cat.# CCL-2 |
| HeLa RIT1 KO-1 (clone 1) | This paper. | N/A |
| HeLa RIT1 KO-2 (clone 2) | This paper. | N/A |
| Human: U2-OS | ATCC | Cat.# HTB-96 |
| Human: HCT-116 | ATCC | Cat.# CCL-247 |
| Human: hTERT RPE-1 | ATCC | Cat.# CRL-4000 |
| hTERT RPE-1 RIT1 KO (three clonal lines) | This paper. | N/A |
| hTERT RPE-1 RIT1 WT (three clonal lines) | This paper. | N/A |
| Oligonucleotides |  |  |
| SMARTpool: ON-TARGETplus RIT1 | Horizon | N/A |
| ON-TARGETplus Non-targeting Control Pool | Horizon | Cat.# D-001810-10-05 |
| Recombinant DNA |  |  |
| psPAX2 | gift from Didier Trono | Addgene plasmid #12260 |
| pMD.2G | gift from Didier Trono | Addgene plasmid #12259 |
| pDONR221 | Invitrogen | Cat.# 12536017 |
| pDEST566 | gift from Dominic Esposito | Addgene plasmid #11517 |
| pDEST527 | gift from Dominic Esposito | Addgene plasmid #11518 |
| pLenti6-H2B-mCherry | (Pemble et al., 2017) | Addgene plasmid #89766 |
| pSpCas9(BB)-2A-GFP (PX458) | (Ran et al., 2013) | Addgene plasmid #48138 |
| PX458-sgRIT1-1 | This paper. | N/A |
| PX458-sgRIT1-2 | This paper. | N/A |
| pGEX-MAD2 | This paper. | N/A |
| pGEX-MAD2 ΔC | This paper. | N/A |
| pGEX-MAD2 LL | This paper. | N/A |
| pGEX-MAD2 V193N | This paper. | N/A |
| pGEX-MAD2 L13A | This paper. | N/A |
| pGEX-MAD2 ΔN15 | This paper. | N/A |
| pGEX-p31<sup>comet</sup> | This paper. | N/A |
| pGEX-RIT1 | This paper. | N/A |
| pGEX-RIT1 (*D. rerio*) | This paper. | N/A |
| pGEX-RIC (*D. melanogaster*) | This paper. | N/A |
| pGEX-RGL3 RBD | This paper. | N/A |
| pGEX-CDC20 111-138 | This paper. | N/A |
| pDEST527-RIT1 | This paper. | N/A |
| pDEST527-MAD2 | This paper. | N/A |
| pDEST527-MAD2 RQ | This paper. | N/A |
| pDEST527-MAD2 (*D. rerio*) | This paper. | N/A |
| pDEST527-MAD2 (*D. melanogaster*) | This paper. | N/A |
| pDEST566-p31<sup>comet</sup> | This paper. | N/A |
| pDEST566-p31<sup>comet</sup> ΔN | This paper. | N/A |
| pDEST27-RIT1 | This paper. | N/A |
| pDEST27-RIT2 | This paper. | N/A |
| pDEST27-RIT1 ΔN | This paper. | N/A |
| pDEST27-RIT1 ΔC | This paper. | N/A |
| pDEST27-RIT1 ΔC5 | This paper. | N/A |
| pDEST27-RIT1 ΔC10 | This paper. | N/A |
| pDEST27-RIT1 ΔC15 | This paper. | N/A |
| pDEST27-RIT1 ΔC20 | This paper. | N/A |
| pDEST27-RIT1 Ct | This paper. | N/A |
| pDEST27-RIT1 GSL | This paper. | N/A |
| pDEST27-RIT1/RIT2 Ct | This paper. | N/A |
| pDEST27-RIT1 A57G | (Castel et al., 2019) | N/A |
| pDEST27-RIT1 A77P | (Castel et al., 2019) | N/A |
| pDEST27-RIT1 F82L | (Castel et al., 2019) | N/A |
| pDEST27-RIT1 M90I | (Castel et al., 2019) | N/A |
| pDEST27-RIT1 S209A | This paper. | N/A |
| pDEST27-RIT1 S209D | This paper. | N/A |
| pDEST27-RIT1 S209E | This paper. | N/A |
| pDEST27-HRAS | (Rodriguez-Viciana et al., 2004) | N/A |
| pDEST27-NRAS | Rodriguez-Viciana et al., 2004) | N/A |
| pDEST27-KRAS | Rodriguez-Viciana et al., 2004) | N/A |
| pDEST27-RRAS | Rodriguez-Viciana et al., 2004) | N/A |
| pDEST27-MRAS | Rodriguez-Viciana et al., 2004) | N/A |
| pDEST27-TC21 | Rodriguez-Viciana et al., 2004) | N/A |
| pDEST27-RAP1 | Rodriguez-Viciana et al., 2004) | N/A |
| pDEST27-RAP2 | Rodriguez-Viciana et al., 2004) | N/A |
| pDEST27-RHEB | Rodriguez-Viciana et al., 2004) | N/A |
| pDEST302-3xFLAG-EV | This paper. | N/A |
| pDEST302-3xFLAG-RIT1 | This paper. | N/A |
| pDEST663-3xFLAG-EV | This paper. | N/A |
| pDEST663-3xFLAG-RIT1 | This paper. | N/A |
| pDEST663-3xFLAG-RIT1 M90I | This paper. | N/A |
| pDEST663-3xFLAG-RIT1 M90I/S209A | This paper. | N/A |
| pDEST663-3xFLAG-RIT1 M90I/S209A | This paper. | N/A |
| pDEST663-3xFLAG-RIT1 M90I/S209A | This paper. | N/A |
| pDEST686-3xFLAG-mNeonGreen RIT1 | This paper. | N/A |
| pCDNA3-HA-MAD2 | This paper. | N/A |
| pCDNA3-HA-MAD2 S295A | This paper. | N/A |
| pCDNA3-HA-MAD2 S295D | This paper. | N/A |
| Software and Algorithms |  |  |
| Prism 8 | GraphPad | https://www.graphpad.com/ |
| ZEN 2.1 Blue | Zeiss | https://www.zeiss.com |
| Fiji | (Schindelin et al., 2012) | https://imagej.net/Fiji |
| Skyline v20 | (Pino et al., 2020) | https://skyline.ms |
| Excel | Microsoft | https://www.microsoft.com |

### RESOURCE AVAILABILITY

#### Lead Contact

Further information and requests for resources and reagents should be directed to and will be fulfilled by the Lead Contact, Frank McCormick.

#### Meterial Availability

All unique/stable reagents generated in this study are available from the Lead Contact with a completed Materials Transfer Agreement.

#### Data and Code Availability

This study did not generate/analyze datasets or code.

### EXPERIMENTAL MODEL AND SUBJECT DETAILS

#### Cells and culture conditions

HEK-293T, HeLa, U2-OS, HCT-116, and hTERT RPE-1 cells were obtained from the American Type Culture Collection (ATCC). HEK-293T, HeLa and U2-OS cells were cultured in Dulbecco’s modified Eagle’s medium (DMEM) supplemented with 10% Fetal Bovine Serum (FBS). hTERT RPE-1 cells were cultured in DMEM/F12 medium supplemented with 10% FBS. Cells were grown in a humidified incubator with 5% CO2 at 37 °C. Validation procedures are as described by the manufacturer. Cell lines were regularly tested and verified to be mycoplasma negative using MycoAlert PLUS Mycoplasma Detection Kit (Lonza).

### METHOD DETAILS

#### Reagents, antibodies and immunoblotting

Nocodazole, Reversine, Dinaciclib, RO-3306, JNK-IN-8, CHIR-99021, TG003, Losmapimod, and Trametinib were purchased from Selleckchem. MG132, Thymidine, ATP, phospho-creatine, GDP and GTPγS were purchased from Sigma-Aldrich. RIT1 (202-216), pS209 RIT1 (202-216), MBP1 (SWYSYPPPQRAV) and control (SYWPQRAPPSVY) peptides were obtained from GenScript. Antibodies against p-ERK (4370; 1:1000), ERK1/2 (4696; 1:2000), p-MEK (9154; 1:1000), MEK1/2 (4694; 1:1000), Cyclin B1 (4135, 1:1000), HA (3724, 1:1000), FLAG (14793; 1:1000), GST (2625; 1:1000), and (K/H)pSP (9477, 1:1000) were obtained from Cell Signal Technology. Antibodies recognizing Securin (PTTG) (sc-56207; 1:250), MAD2 (sc-47747; 1:250) and GST (sc-138; 1:1000) were obtained from Santa Cruz Biotechnology. RIT1 antibody (ab53720; 1:1000) was from Abcam. p31^comet^ antibody (MABE451; 1:500) was from EMD Millipore. MAD2 antibody (A300-301A; 1:1000) was from Bethyl Laboratories. βActin (A2228; 1:10000), α-Tubulin (T6199; 1:5000), FLAG (F1804; 1:2000) and His_6_ (05-949; 1:1000) antibodies were purchased from Sigma-Aldrich. Whole cell lysates were prepared using RIPA Buffer (50 mM Tris-HCl (pH 8.0), 150 mM NaCl, 0.5% Deoxycholate, 0.1% SDS, 1% IGEPAL CA-630) supplemented with protease and phosphatase inhibitor cocktails (Sigma-Aldrich). 20-30 μg of total protein was loaded per well of pre-casted NuPAGE gels (Life Technologies). For immunoblot detection, samples were separated by SDS-PAGE and transferred onto nitrocellulose membranes. Membranes were blocked using 5% skimmed milk in TBST buffer for 1 hour and incubated with appropriate primary antibodies overnight. Detection was performed using secondary antibodies conjugated to DyLight680 (611-144-002; 1:10,000) or DyLight800 (610-145-002; 1:10,000) (Rockland), and visualized with a LI-COR Odyssey infrared scanner.

#### Plasmids, cloning and transfection

All RIT1 mutations used in this paper were generated by standard PCR-based mutagenesis in the pDONR223-RIT1 template. These included ΔN, ΔC (and all C-terminal truncations), GSL, S209A, S209D, S209E, A57G, A77P, F82L, and M90I. Mutagenesis primer sequences are available upon request. RIT2 (Isoform 1, NM_002930.4), RIT1 (RIT2 Ct) chimera, and mNeonGreen-RIT1 were synthesized as a gene block and cloned into pDONR221 (Invitrogen) using BP reaction. LR Gateway cloning yielded mammalian expression vectors with indicated tags. For N-terminal GST-tagged proteins, entry clones were cloned into pDEST27 destination vector (Invitrogen). For FLAG-tagged proteins, entry clones were cloned by multisite gateway cloning into pDEST302, pDEST663 or pDEST686 (a gift from Dominic Esposito, Frederick National Lab), and designed to express N-terminal 3xFLAG tag fusion proteins driven by an EF1a promoter (Wall et al., 2014). Empty vector (EV) plasmid controls were generated using a gateway recombination cassette containing a stop codon followed by an untranslated stuffer sequence. The GST-tagged Ras Family GTPases panel was cloned in the pDEST27 vector and was previously described (Rodriguez-Viciana et al., 2004). MAD2 cDNA (NM_002358.3) was purchased from GeneCopoeia as a Gateway entry clone and was recombined into pcDNA3-HA destination vector to be expressed an N-terminal HA-tagged fusion protein. MAD2 S195A and S195D were generated by PCR-based mutagenesis. All plasmid transfections in this study were performed using JetPRIME transfection reagent (Polyplus Transfection) according to manufacturer’s instructions.

Constructs used for bacterial expression were generated from as follows: a gene block containing *E. coli* codon-optimized human RIT1 (a.a.1-219), with an N-terminal TEV cleavage site followed by a FLAG-tag, was cloned into pDONR221 (Invitrogen). Similarly, gene blocks encoding *E. coli* codon-optimized human MAD2 (1-205) and p31^comet^ (1-274), both with an N-terminal TEV cleavage site, were synthesized and cloned into pDONR221. MAD2 R133E/Q134A (RQ); ΔC (1-195); Loop-less (LL), in which residues 109-117 are replaced with a Gly-Ser-Gly linker; V193N; L13A; and ΔN15 (16-195); and p31^comet^ΔN were generated by standard PCR-based mutagenesis(Mapelli et al., 2007). p31^comet^ΔN (50-274) was generated to enhance protein stability through deletion of its non-conserved and disordered N-terminal fragment (Yang et al., 2007). Zebrafish *(Danio rerio)* and Fruit fly *(Drosophila melanogaster)* MAD2 and RIT1 orthologs were synthesized as *E. coli* codon-optimized gene blocks and cloned into pDONR221. RGL3-RBD (604-703) was synthesized as an *E. coli* codon-optimized gene block and subcloned into pGEX-6P-3 at EcoRI and XhoI restriction sites. All plasmids were verified by Sanger sequencing.

#### RNA interference

The short interfering RNAs (siRNAs) used in this study are siRIT1 (SMARTpool: ON-TARGETplus RIT1, Horizon) and siNC (ON-TARGETplus Non-targeting Control Pool, D-001810-10-05, Horizon). Cells were transfected with siRNAs using Lipofectamine RNAi Max Transfection Reagent (Life Technologies) according to manufacturer’s instructions.

#### Generation of CRISPR/Cas9-mediated knockout cells

RIT1 knockout (KO) clones were generated using two sgRNA targeting exon 2 of RIT1: sgRIT1-1: GATTCTGGAACTCGCCCAGT and sgRIT1-2: GGAGTACAAACTAGTGATGC. Briefly, Parental cells were transiently transfected with plasmid encoding SpCas9, EGFP (PX458, Addgene, plasmid #48138) and sgRIT1-1 or sgRIT1-2. 48 h post-transfection, GFP+ cells were single-cell sorted into 96-well plates using a SONY SH800 FACS. Clones were expanded and KO clones were validated by Sanger sequencing and Western blot analysis.

#### Lentiviral transduction

Lentivirus was produced by co-transfection of HEK-293T cells with a lentiviral vector and the packaging plasmids psPAX2 (Addgene, plasmid #12260) and pMD.2G (Addgene, plasmid #12259) at a ratio of 1.25:1.0:0.25. The supernatant was collected 72 hours post-transfection and filtered through a 0.45 μm filter. Cells were transduced with lentiviral-containing supernatant supplemented with 0.8 μg/ml polybrene (Sigma-Aldrich). Stably transduced cells were selected with appropriate antibiotic and maintained in media containing 50% antibiotic used during selection.

#### Bacterial protein expression and purification

Full-length recombinant RIT1 protein was obtained by gateway cloning pDONR-TEV-FLAG-RIT1 WT or S209A into pDEST566 (Addgene, plasmid #11517) containing an N-terminal hexahistidine-maltose binding protein (His_6_-MBP) tag. Expression constructs were transformed into the *E. coli* strain BL21(DE3) (New England Biolabs). Protein expression was induced in cultures at OD_600_ between 0.4-0.6 with 200 μM Isopropyl β-D-1-thiogalactopyranoside (IPTG) (Sigma-Aldrich) for 14-16 h at 18°C. Cells were lysed by sonication in 100 mM Sodium phosphate (pH 6.0), 300 mM NaCl, 10 mM MgCl2, 5% glycerol, 2 mM DTT, 1 mg/ml DNAse I, 0.2 mg/ml lysozyme, 30 mM imidazole and protease inhibitor cocktail (Sigma-Aldrich, P8849). After clearing, the lysate was loaded on a HisTrap FF metal chelating column (Cytiva Life Sciences) equilibrated in 100 mM Sodium phosphate (pH 6.0), 300 mM NaCl, 10 mM MgCl2, 5% glycerol, 30 mM imidazole. Bound proteins were eluted with 300 mM imidazole. Fractions containing RIT1 were pooled and dialyzed overnight in the presence of TEV-protease (Sigma-Aldrich). Cleaved protein was recovered by subtractive purification, concentrated by ultrafiltration, and further separated by size exclusion chromatography (SEC) on a Superdex-75 column (Cytiva Life Sciences) equilibrated in 100 mM Sodium phosphate (pH 6.0), 300 mM NaCl, 10 mM MgCl2, 5% glycerol, 0.5 mM TCEP. RIT1 containing fractions were pooled, concentrated and frozen in liquid nitrogen. The entire purification scheme was carried out at 4°C. Recombinant p31^comet^ WT and ΔN protein were expressed and purified analogously but with the use of 50 mM Tris-HCl (pH 8.0) buffer without MgCl2.

Recombinant MAD2 WT and RQ were expressed from pDEST527 (Addgene, plasmid #11518) containing an N-terminal hexahistidine-tag in BL21(DE3) following the same conditions described above in 50 mM Tris-HCl (pH 8.0) buffer without MgCl2. After subtractive purification, TEV-cleaved MAD2 protein was dialyzed overnight in anion-exchange (AE) buffer (10 mM Tris-HCl (pH 8.0), 30 mM NaCl, 5% glycerol and 1 mM DTT). MAD2 was loaded onto an AE Resource-Q column (Cytiva Life Sciences) equilibrated in AE Buffer. The protein was eluted using a NaCl gradient, concentrated and flash-frozen in liquid nitrogen.

Recombinant GST-tagged proteins were expressed from a pGEX-6 plasmid transformed into BL21 (DE3) cells. Expression was induced with 0.2 mM IPTG for 14-16 h at 18°C. Cells were lysed by sonication in 50 mM Tris-HCl (pH 8.0), 300 mM NaCl, 5% glycerol, 1 mM DTT. Proteins were immobilized on Glutathione sepharose 4B beads (Cytiva Life Sciences), washed extensively, and stored as a 50% glycerol bead suspension at −20°C.

#### Immunoprecipitation and GST pulldown assays

GST pulldown assays with recombinant proteins were performed by diluting indicated proteins in 500 μl of pulldown buffer (10 mM Tris-HCl (pH 8.0), 150 mM NaCl, 0.1% IGEPAL CA-630, 10% glycerol) and 20 μl of Glutathione sepharose 4B beads (Cytiva Life Sciences) for 1 h at 4°C with end over end rotation. Beads were rinsed three times with pulldown buffer and resuspended in LDS sample buffer (Life Science Technologies) for immunoblot or Coomassie staining of SDS-PAGE gels. For nucleotide loading, RIT1 was incubated in 100 μl GTPase loading buffer (20 mM Tris 7.5, 25 mM NaCl, 5 mM EDTA) containing 2 mM GTP or GTPγS, for 30 min at 30°C. Samples were chilled on ice and MgCl2 was added to a final concentration of 10 mM. RGL3-RBD was used as a positive control for nucleotide loading due to the GTP-dependent interaction between RIT1 and RGL3 (Shao and Andres, 2000).

For GST pulldown of proteins from cell lysates, 3 x 10^6^ HEK-293T cells were transfected with 4 μg total DNA of indicated plasmids. 24 hours after transfection, cells were rinsed with ice-cold PBS and lysed with 1 ml of Lysis buffer (50 mM Tris-HCl (pH 8.0), 150 mM NaCl, 1% IGEPAL CA-630, 10% glycerol) supplemented with protease and phosphatase inhibitor cocktails (Sigma-Aldrich). Lysates were cleared by centrifugation and incubated with 20 μl of Glutathione Sepharose 4B beads for 4 h at 4°C with end over end rotation. Beads were rinsed three times with Lysis buffer and resuspended in LDS sample buffer.

For immunoprecipitation of FLAG-RIT1 from cell cycle synchronized HeLa cells, approximately 10^6^ cells were rinsed with PBS and harvested using a cell scraper at indicated timepoints, spun down and frozen. Cells were then lysed with RIPA Buffer (50 mM Tris-HCl (pH 8.0), 150 mM NaCl, 0.5% Deoxycholate, 0.1% SDS, 1% IGEPAL CA-630) supplemented with protease and phosphatase inhibitor cocktails (Sigma-Aldrich), cleared by centrifugation and incubated with 20 μl anti-FLAG M2 agarose beads (EMD Millipore) for 4 hours at 4°C with end over end rotation. Beads were rinsed three times with RIPA buffer and resuspended in LDS sample buffer.

#### Cell cycle synchronization

Synchronization of cells at G1/S boundary was performed with a double thymidine block. Briefly, ~50% confluent cells were treated with 2 mM thymidine (Sigma-Aldrich) for 20 h, rinsed twice and released into drug-free media for 9 h, then treated again with 2 mM thymidine for 18 h. Synchronization of cells in prometaphase was done by addition of 100 ng/ml nocodazole (Selleckchem) 4 h after release from a single-thymidine block and incubated for 10 h. Mitotic cells were collected by mechanical shake off.

#### Subcellular protein fractionation

For subcellular fractionation of endogenous RIT1, Asynchronous cells were harvested with trypsin-EDTA and then centrifuged at 500 x g for 5 min. Nocodazole-arrested mitotic cells were harvested by mitotic shake-off and centrifuged at 500 x g for 5 min. Cell pellets were rinsed with ice-cold PBS and transferred to 1.5 ml tubes and pelleted by centrifugation at 500 x g for 5 min. Cell pellets were then lysed and proteins were fractionated using the Subcellular Protein Fractionation Kit for Cultured Cells (Thermo Scientific) according to manufacturer’s instructions.

#### Size Exclusion Chromatography

Equal molar ratios of indicated proteins were incubated at 10 μM final concentration in PBS (pH 7.4), 0.5 mM TCEP for 1 hour at 25°C. Samples were chilled on ice and centrifuged (15,000 rpm) to remove any precipitates before loading onto a Superdex 200 3.2/300 column equilibrated in PBS (pH 7.4), 0.5 mM TCEP. All samples were eluted under isocratic conditions at 4°C with a flow rate of 0.035 ml min^−1^. Elution profiles were monitored at 280nm. Elution fractions were separated by SDS-PAGE and stained with Coomassie.

#### Microscopy/Metaphase spreads

For analysis of mitotic errors (chromosome segregation errors), HCT-116 cells were grown on #1.5 coverslips, rinsed with PBS and fixed with 100% methanol. Coverslips were quickly hydrated and mounted with Prolong Gold Antifade mounting media with DAPI (Invitrogen). Cells were imaged on a Zeiss AxioImager M1 fluorescent microscope equipped with a 40x/0.75 Plan-Neofluar objective (Zeiss) and controlled with ZEN imaging software (Zeiss). For each biological replicate, at least 60 anaphase cells were analyzed per condition.

For metaphase chromosome spread analysis, cells were treated with 0.1 μg/ml Colcemid (Thermo Fisher Scientific) for 2 h, trypsinized and spun down. Cell pellets were gently resuspended in 2 ml hypotonic solution (75 mM KCl) added dropwise while mixing cell suspension, followed by 15 min incubation at 37°C. Cells were spun down again and resuspended in 5 ml of Carnoy’s fixative (3:1, methanol:glacial acetic acid, made fresh) added dropwise to cells. Cells were allowed to fix at room temperature for 20 min, then centrifuged and rinsed twice with Carnoy’s fixative. Cells were dropped onto clean coverslips and rinsed with fixative to remove debris. Coverslips were placed in a humidity chamber for 10 min, then allowed to air dry for 24-72 h. Chromosome spreads were mounted with ProLong Gold Antifade mounting medium with DAPI (Thermo Fisher Scientific) and imaged on a Zeiss AxioImager M1 fluorescent microscope equipped with a 63x/1.25 Plan-Neofluar oil objective (Zeiss). Images of at least 75 chromosome spreads were captured per condition and experimenter blinded before being quantified using Fiji (Schindelin et al., 2012).

#### Live cell imaging

For mitotic duration experiments, cells were engineered to stably express Histone H2B-mCherry (pLenti6-H2B-mCherry, Addgene, plasmid #89766), and seeded onto 12-well #1.5 glass bottom plates (Cellvis). Drug treatments were performed 1 h before imaging. Time-lapse images were captured on a Nikon Ti-E inverted wide-field fluorescent microscope equipped with a 20x/0.75 Plan Apo air objective (Nikon). HeLa and U2-OS cells were imaged at 5 min intervals, hTERT-RPE1 cells at 2 min intervals, for 20 hours. The microscope was equipped with an incubation chamber (Okolab), providing a humidified atmosphere at 37 °C with 5% CO2. Mitotic length was quantified as the duration between nuclear disassembly and anaphase onset. Mitotic error rates in hTERT-RPE1 cells were determined from live-cell images used to assess mitotic length. Images were analyzed using Fiji.

For RIT1 subcellular localization experiments, hTERT-RPE1 cells stably express Histone H2B-mCherry and mNeonGreen (mNG)-RIT1 were seeded on seeded onto 12-well #1.5 glass bottom plates and allowed to adhere for 24-48 h. Prior to imaging, cells were exchanged into imaging media: FluoroBrite DMEM (Thermo Fisher Scientific) supplemented with 10% FBS and 4 mM GlutaMAX (Thermo Fisher Scientific). Images were acquired as a series of 0.9 μm z-stacks with a Plan Apo 40x/0.95 Corr (DIC N2 / 40X I) 227.5 nm/pixel objective (Nikon) at 5 min intervals on a Nikon Ti-E inverted CSU-22 spinning disk confocal microscope equipped with an incubation chamber (Okolab), providing a humidified atmosphere at 37 °C with 5% CO2. Images were analyzed using Fiji. For quantitative analysis, a single central plane along the z-axis was used. For PM/Cyto. ratio calculations, semi-automated recognition of the cell boundary within the Fiji program was used to create regions of interests designating the plasma membrane and cytoplasm, and mean fluorescence intensity was calculated within each region.

#### Kinase Assays

For phosphorylation of RIT1 using mitotic cell extract, metaphase arrested HEK-293T cells were harvested, rinsed with PBS and lysed with 4x pellet volume of Lysis buffer (50 mM Tris-HCl (pH 8.0), 150 mM NaCl, 1% IGEPAL CA-630, 10% glycerol) supplemented with 10 μM MG132 (Sigma-Aldrich) protease and phosphatase inhibitor cocktails (Sigma-Aldrich). The lysate was cleared by centrifugation for 10 min at 15,000 rpm. 1 ml of cleared lysate was incubated with indicated drugs (10 μM final concentration) or DMSO for 30 min at 4°C. No lysate control consisted of 1 ml Lysis buffer. Bacteria purified GST-RIT1 or GST protein bound to sepharose beads was added to lysates at 0.5 μM final concentration, together with 10 mM MgCl2 and 0.1 mM ATP. Reactions were incubated at 30°C for 1 h with end over end rotation. Tubes were chilled on ice for 5 min, beads were then centrifuged and rinsed three times with RIPA buffer and resuspended in LDS sample buffer.

CDK1/Cyclin B1 kinase assays were conducted using recombinant active CDK1/Cyclin B1 purchased from SignalChem (C22-10G). 20 μl reactions containing 200 ng CDK1/Cyclin B1 protein and 3 μg of RIT1 protein diluted in Kinase assay buffer (5 mM MOPS, pH7.2, 2.5 mM β-glycerol-phosphate, 10 mM MgCl2, 1 mM EGTA, 0.4 mM EDTA, 50 ng/μl BSA) with or without 3mM ATP were incubated at 30°C for 1 h. Reactions were stopped by addition of LDS sample buffer for SDS-PAGE or flash-frozen in liquid nitrogen for mass spectrometry analysis.

#### Mass Spectrometry

For identification of RIT1 binding partners, approximately 2 x 10^7^ HEK-293T cells were transiently transfected with 8 μg of plasmid (FLAG-RIT1 or EV control) and immunoprecipitated as described above with buffer containing 50 mM Tris-HCl (pH 8.0), 150 mM NaCl, 1% IGEPAL CA-630, 10% glycerol and supplemented with protease and phosphatase inhibitor cocktails (Sigma-Aldrich). Magnetized beads were washed with ice-cold 20 mM Tris-HCl (pH 8.0), 2 mM CaCl2 buffer and frozen prior to trypsin digest.

Protein pulldown samples were on-bead digested with trypsin as previously described (Castel et al., 2019). Briefly, the beads were resuspended in 9 μL of 20 mM Tris-HCl pH 8.0. The proteins were reduced with DTT, 5 mM final concentration, at room temperature for 30 min; alkylated with iodoacetamide, 15 mM final concentration, at room temperature for 10 min; and digested with 500 ng of trypsin (Sigma Trypsin Singles, T7575) at 37°C overnight. In vitro kinase assay samples, 20 μL each, were digested using the same protocol. All samples were desalted with ZipTip u-C18 pipette tips (Millipore), vacuum dried, and reconstituted in 15 μL of 0.1% formic acid for analysis by LC-MS/MS.

LC-MS/MS was carried out on Acquity UPLC M-Class system (Waters) online with Orbitrap Fusion Lumos Tribrid Mass Spectrometer (Thermo Fisher Scientific). Reversed-phase chromatography was performed on a 15 cm silica-C18 EasySpray column (Thermo Fisher Scientific) at 45°C with a binary buffer system (Buffer A = 0.1% formic acid in water; Buffer B = 0.1% formic acid in acetonitrile) and a flow rate of 400 nL/min. The sample was loaded at 2% B for 20 min followed by a 2-60% B gradient over 60 min, followed by a brief wash at 80% B and equilibration at 2% B. The mass spectrometer was operated in Full-MS/ddMS2 mode with one survey scan (375-1500 m/z, R=120,000, AGC target of 4e5), followed by a series of data-dependent HCD MS2 scans not to exceed a 3 sec cycle (AGC target of 5e4, max IT 100 ms, R=30,000, isolation window 1.6 m/z, NCE 30%, stepped collision 5%, and 30 s dynamic exclusion).

MS raw data files were converted to peak list files using Proteome Discoverer v. 1.4 (Thermo Fisher Scientific) and searched using Protein Prospector (Baker et al., 2011; Chalkley and Baker, 2017) version 6.0.0 against human SwissProt database (UniProt Consortium, 2019) downloaded on 01/08/2018 (or a subset of this database with RIT1, CDK1 and CCNB1 entries only when searching in vitro kinase assay samples) and a corresponding random concatenated decoy database. Other settings included the default “ESI-Q-high-res” parameters with trypsin as the protease, up to two allowed missed cleavage sites, Carbamidomethyl-C as a constant modification, default variable modifications for pulldown samples (or default variable modifications plus phosphorylation at STY for in vitro kinase assay samples), up to 3 modifications per peptide, and 5 ppm precursor mass and 15 ppm fragment mass tolerance. False discovery rate of <1% was used as the cutoff for peptide expectation values. Protein Prospector search results were exported in BiblioSpec format compatible with downstream analysis in Skyline (Pino et al., 2020). Quantitation of peptide and protein abundances was carried out in Skyline v20 by quantifying MS1 precursor peak areas with normalization by median centering (Schilling et al., 2012). Peptides shared by multiple proteins in the database were excluded.

#### APC/C Ubiquitination Assay

Detection of APC/C activity using mitotic checkpoint active cell extracts was previously described (Braunstein et al., 2007). HeLa RIT1 KO cells arrested in prometaphase were harvested, rinsed with icecold PBS and resuspended in 75% pellet volume of hypotonic buffer (20 mM HEPES, pH 7.6, 5 mM KCl, 1 mM DTT) containing protease inhibitor cocktail (Roche). Cells were lysed with multiple rounds of freeze-thawing and were cleared by centrifugation at 13,000 rpm for 1 h at 4°C. Cleared lysate was supplemented with glycerol to 10% (v/v), aliquoted and frozen in liquid nitrogen. The protein concentration of cell extract was ~20 mg/ml. APC/C activity reactions were carried in samples containing 50% (v/v) mitotic cell extract diluted in buffer with final concentrations of the following: 10 mM Tris-HCl, pH 7.6, 5 mM MgCl2, 1 mM DTT, 1 mg/ml Ubiquitin (Boston Biochem; U-100H), 10 mM phosphocreatine, 0.5 mM ATP, 10 μg/ml UbcH10 (Boston Biochem; E2-650), and 50 μg/ml Creatine phosphokinase (Sigma-Aldrich). Recombinant RIT1 protein was buffer exchanged into assay buffer containing 20 mM HEPES pH 7.4, 300 mM NaCl, 1 mM TCEP, 5% glycerol before being added to samples. Control conditions received equal volumes of assay buffer. Samples were incubated on ice for 1 h, then transferred to 30°C. 4 μl samples were withdrawn at indicated times and rapidly quenched with LDS sample buffer. Degradation of Cyclin B1 and Securin was followed by immunoblotting and was normalized to MAD2 protein levels. Densitometry analysis was performed using Fiji.

#### Statistical Analysis

Statistical analysis was performed using GraphPad Prism 7.0 (GraphPad). Results are expressed as mean ± s.e.m. For each scatterplot, the horizontal line represents the mean. No statistical methods were used to predetermine the sample size. Experiments analyzed by immunoblotting were repeated 2-4 times with similar results. For chromosome spread analysis, investigators were blinded to sample allocation.

---

**Figure S1.**
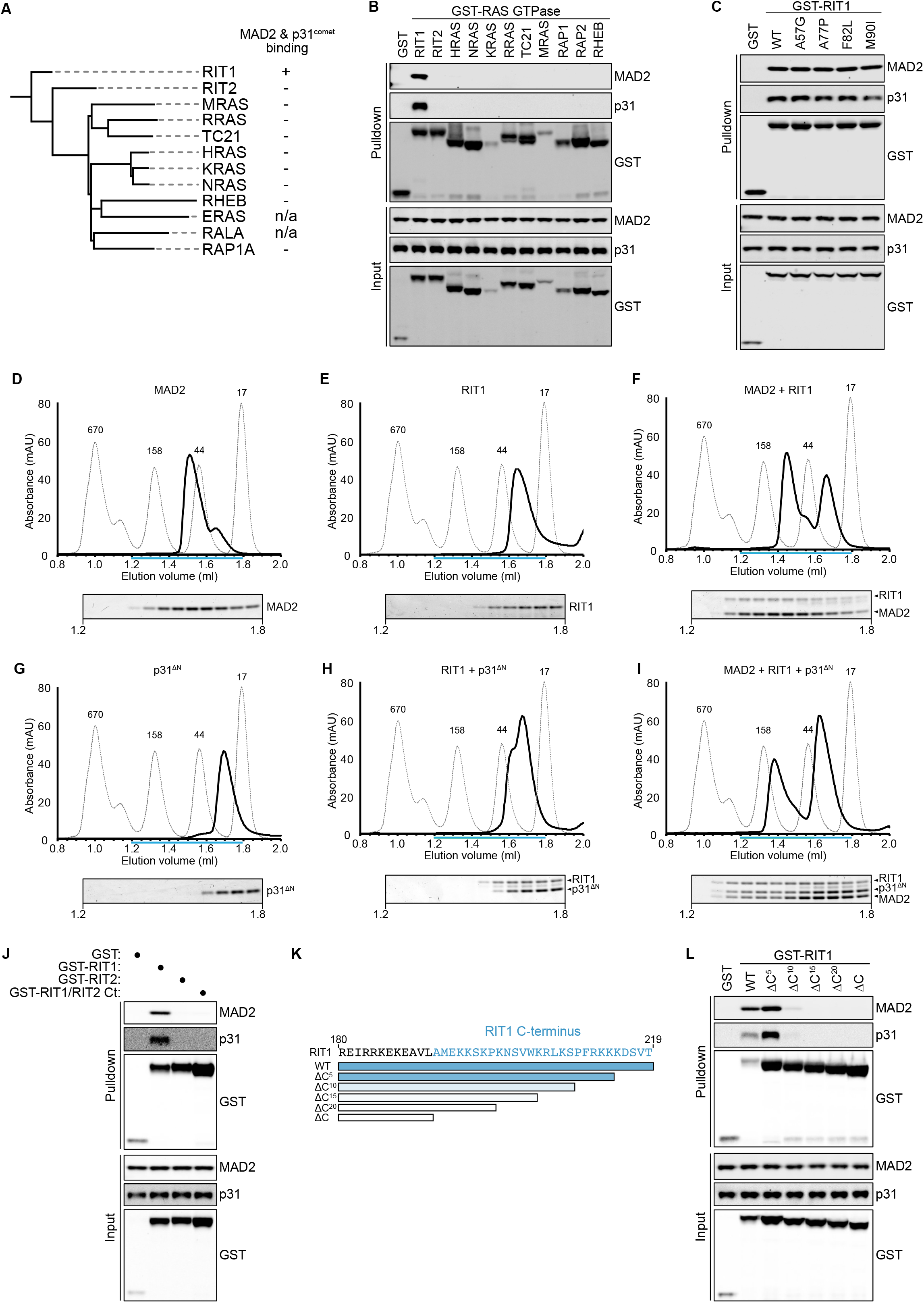
RIT1, but not other Ras GTPases, forms a complex with MAD2 and p31^comet^ mediated by its C-terminal extension, related to Figure 1. (**A**) Summary of MAD2 and p31^comet^ binding exhibited by closely related Ras family GTPases. (**B**) Representative immunoblot of data summarized in (A) showing proteins precipitated from extracts of HEK-293T cells transfected with GST or a GST-Ras GTPase construct probed for endogenous MAD2 or p31^comet^. (**C**) Analysis of MAD2 and p31^comet^ binding to disease-associated RIT1 mutants. Immunoblots of proteins precipitated from extracts of HEK-293T cells transfected with GST or indicated GST-RIT1 constructs and probed for endogenous MAD2 or p31^comet^. (**D-I**) RIT1, MAD2, and p31^comet^ proteins were allowed to complex at 25°C for 1 hour prior to gel filtration. Elution profiles were compared to that of known standards (thin dashed line). For every chromatogram, the contents of 12 consecutive 50 μ1 fractions eluting between 1.2 ml to 1.8 ml were separated by SDS-PAGE and stained with Coomassie. (**D**) elution profile of bacterially expressed MAD2, with the minor peak corresponding to monomeric MAD2 and the major peak representing dimerization of MAD2 between O-MAD2 and C-MAD2 conformers. (**E**) Elution profile of full-length RIT1, showing a single monomeric peak. (**F**) Elution profile of MAD2 and RIT1 demonstrates complex formation between RIT1 and dimeric MAD2. (**G**) Elution of profile of p31^comet^ ΔN. (**H**) No complex formation between RIT1 and p31^comet^ ΔN can be observed by size exclusion under assay conditions. Identical results are observed with full-length p31^comet^ protein. (**I**) Elution profile RIT1, MAD2, and p31^comet^ shows that incubation of all three proteins allows for the formation of a high molecular weight complex that includes MAD2, RIT1 and p31^comet^. (**J, L**) Immunoblots of proteins precipitated from extracts of HEK-293T cells transfected with GST or indicated GST-RIT1 or RIT2 constructs and probed for endogenous MAD2 or p31^comet^. (**J**) RIT1/RIT2 Ct, a chimeric protein with RIT1 residues 194-219 replaced with RIT2 residues 192-217. (**K**) Diagram of RIT1 C-terminal truncation mutants used in (L).

**Figure S2.**
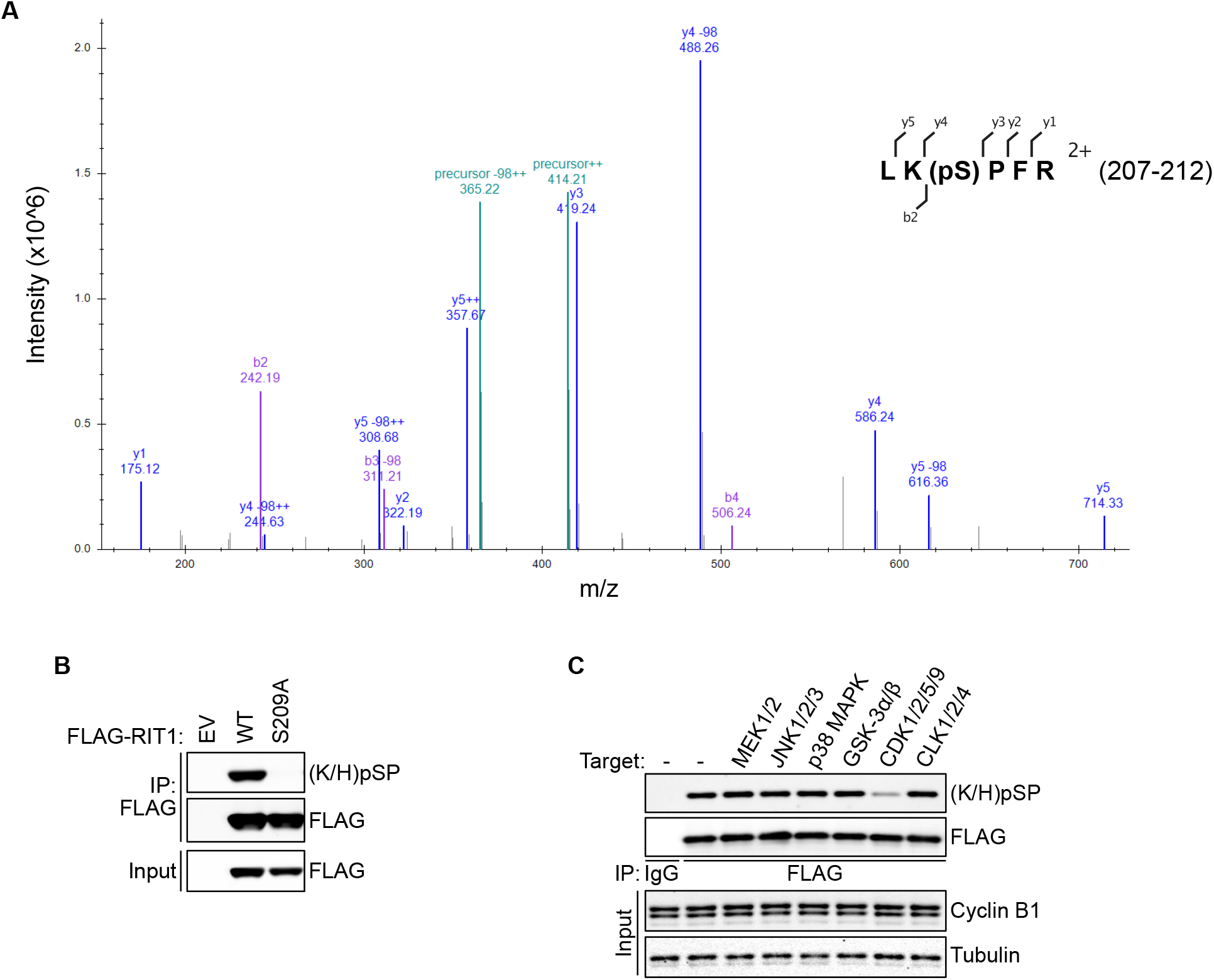
RIT1 is phosphorylated at S209, related to Figure 2. (**A**) Representative MS2 spectrum from RIT1 phosphopeptide. The annotated peptide sequence includes b-series (_⌞_) and y-series (^⌝^) ions, precursor charge state, and the residues within the RIT1 protein sequence. (**B**) Validation of phospho-serine −1(K/H)/+1P antibody for detection of RIT1 S209 phosphorylation. Immunoblot of FLAG-RIT1 immunoprecipitated from HEK-293T cells expressing RIT1 WT or S209A and probed with an antibody against (K/H)pSP. EV, empty vector. (**C**) HeLa cells stably expressing FLAG-RIT1 were arrested in prometaphase and were kept in mitosis with MG132 (10 μM) while being treated with 1 μM of the following inhibitors for 4 hours: Trametinib (MEK1/2i), JNK-IN-8 (JNK1/2/3i), Losmapimod (p38 MAPKi), CHIR-99021 (GSK-3α/βi), Dinaciclib (CDK1/2/5/9i), TG003 (CLK1/2/4i). FLAG-RIT1 was immunoprecipitated and subjected to immunoblotting for detection of S209 phosphorylation.

**Figure S3.**
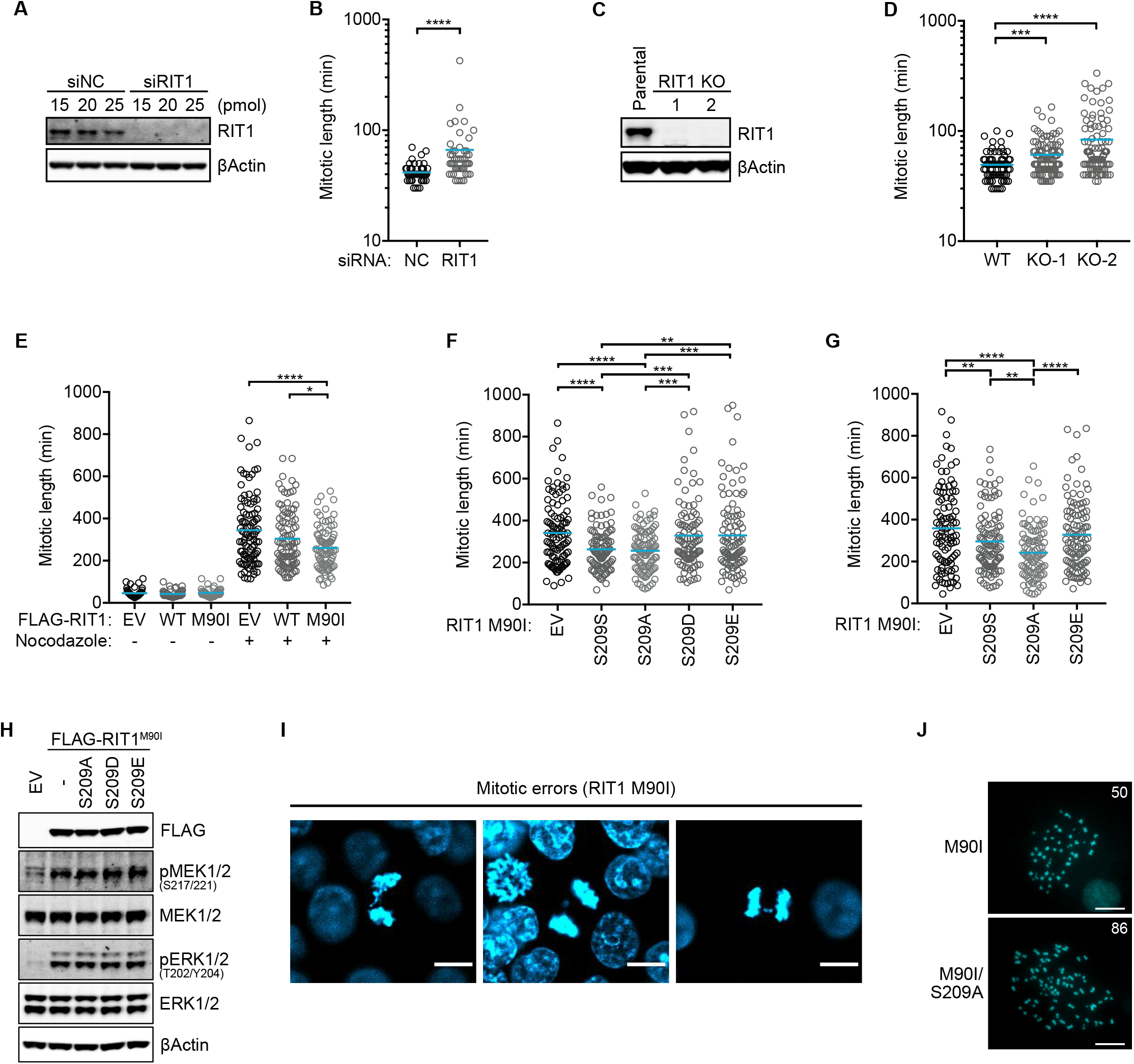
Inhibition of the SAC by RIT1 is dependent on MAD2 and p31^comet^ binding, related to Figure 3. (**A**) Immunoblots of lysates from HeLa cells harvested 72 h after transfection with RIT1 or nontargeting control (siNC) siRNA. (**B**) Duration of mitotic length (NEBD - anaphase onset) assessed by time-lapse microscopy in HeLa cells transfected with indicated siRNA 72 hours prior to the start of imaging. *(n*= 50). (**C**) Immunoblots of lysates from parental HeLa cells and RIT1 KO HeLa clonal cell lines. (**D**) Mitotic length measured as in (B) in Parental HeLa cells (*n*= 75) and in two RIT1 KO clones (*n*= 100). (**E**) Duration of mitotic length as in (B) in HeLa cells stably expressing indicated proteins treated with 15 ng/ml nocodazole (EV, WT, M90I, *n*= 98, 99, 84) or DMSO control (*n*= 50). (**F**, **G**) Duration of mitotic length as in (B) in (F) HeLa, *n =* 100, or (G) hTERT-RPE1, *n* ≥ 100, cells stably expressing indicated proteins treated with 15 ng/ml nocodazole. (**H**) Immunoblots of cell lysates from HEK-293T cells transfected with indicated constructs and serum-starved for 16 hours. EV, empty vector. (**I**) Representative images of mitotic errors from HCT-116 cells stably expressing RIT1 M90I and quantified in Figure 3F. Scale bar, 10 μm. (**J**) Representative images of metaphase spreads quantified in Figure 3G. The inscribed number indicates the chromosome count. Scale bar, 10 μm. (**B, D-G**) *n* indicates the number of cells counted per condition. (**A-F)**Two-sided Student’s *t*-test, bars indicate mean. *\*P*≤ 0.05, *\*\*P*≤ 0.01, *\*\*\*P*≤ 0.001, *\*\*\*\*P*≤ 0.0001.

**Figure S4.**
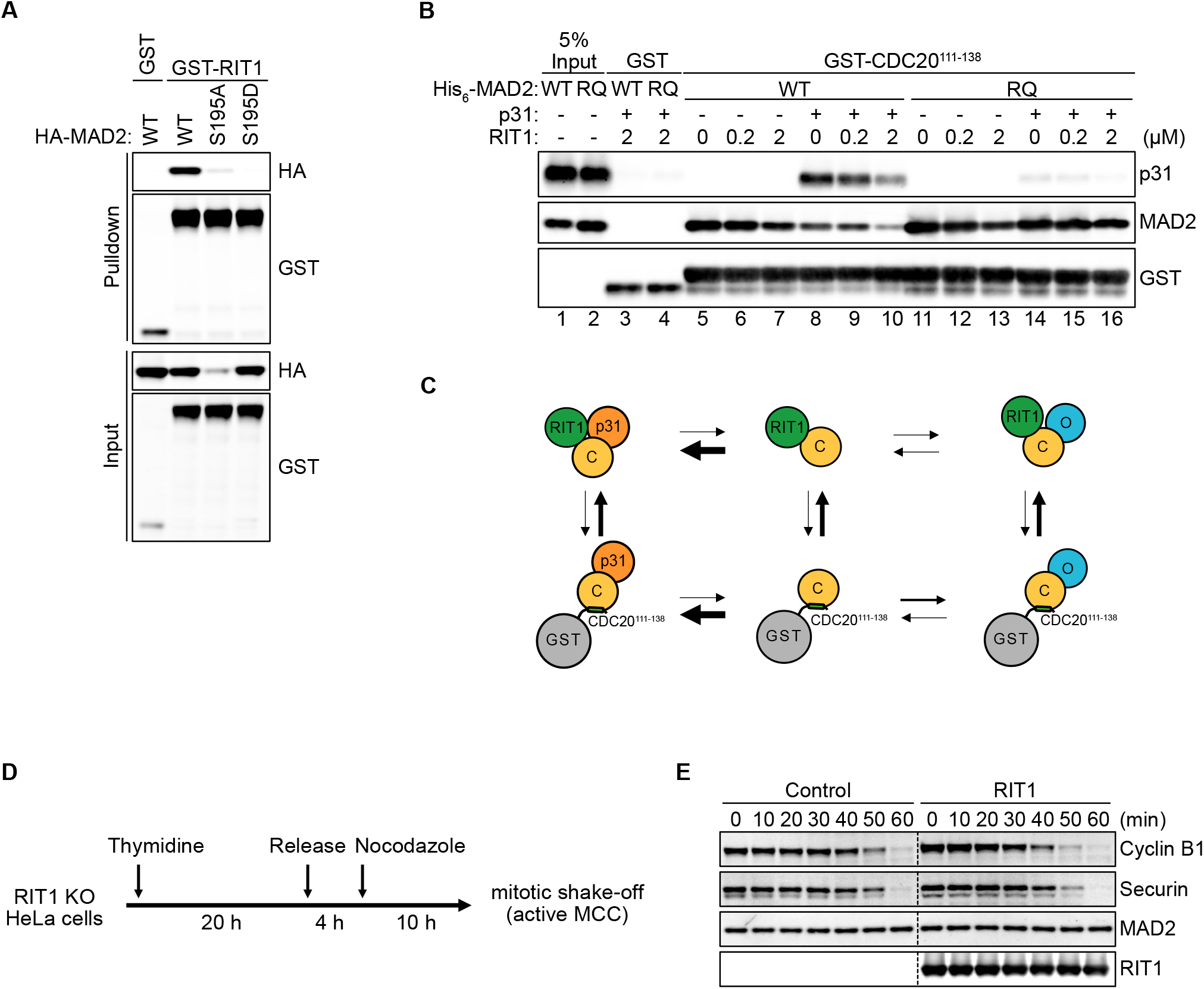
RIT1 inhibits the SAC by cooperating with p31^comet^ to sequester MAD2 away from CDC20, related to Figure 4. (**A**) Immunoblots of proteins precipitated from extracts of HEK-293T cells transfected with GST or GST-RIT1 and hemagglutinin (HA)-tagged MAD2 S195 mutants. S209A mutation significantly reduced expression levels of MAD2, but S209A protein can still be detected in pulldown fractions. (**B**) Immunoblots of GST pulldown assay with 0.5 μM recombinant GST or GST-CDC20 111-138 incubated with or without 0.5 μM p31^comet^, 0.2 μM MAD2 WT or MAD2 R133E/Q134A (RQ) protein and RIT1 protein at indicated concentrations. (**C**) Schematic representation of results from experiment in (A). (**D**) Diagram of cell synchronization strategy to generate mitotic cell extracts with active MCC for APC/C activity assays. (**E**) Representative immunoblots data quantified in Figure 4D. APC/C activity assay following degradation of Cyclin B1 and Securin at indicated time points in the absence of MG132.

## Notes

### Competing Interest Statement

The authors have declared no competing interest.

## REFERENCES

Aoki, Y., Niihori, T., Banjo, T., Okamoto, N., Mizuno, S., Kurosawa, K., Ogata, T., Takada, F., Yano, M., Ando, T., et al. (2013). Gain-of-function mutations in RIT1 cause Noonan syndrome, a RAS/MAPK pathway syndrome. Am. J. Hum. Genet. 93, 173–180.

Baker, P.R., Trinidad, J.C., and Chalkley, R.J. (2011). Modification site localization scoring integrated into a search engine. Mol. Cell. Proteomics MCP 10, M111.008078.

Ben-David, U., and Amon, A. (2020). Context is everything: aneuploidy in cancer. Nat. Rev. Genet. 21, 44–62.

Berger, A.H., Imielinski, M., Duke, F., Wala, J., Kaplan, N., Shi, G.-X., Andres, D.A., and Meyerson, M. (2014). Oncogenic RIT1 mutations in lung adenocarcinoma. Oncogene 33, 4418–4423.

Braunstein, I., Miniowitz, S., Moshe, Y., and Hershko, A. (2007). Inhibitory factors associated with anaphase-promoting complex/cylosome in mitotic checkpoint. Proc. Natl. Acad. Sci. U. S. A. 104, 4870–4875.

Castel, P., Cheng, A., Cuevas-Navarro, A., Everman, D.B., Papageorge, A.G., Simanshu, D.K., Tankka, A., Galeas, J., Urisman, A., and McCormick, F. (2019). RIT1 oncoproteins escape LZTR1-mediated proteolysis. Science 363, 1226–1230.

Chalkley, R.J., and Baker, P.R. (2017). Use of a glycosylation site database to improve glycopeptide identification from complex mixtures. Anal. Bioanal. Chem. 409, 571–577.

Chang, L., and Barford, D. (2014). Insights into the anaphase-promoting complex: a molecular machine that regulates mitosis. Curr. Opin. Struct. Biol. 29, 1–9.

Chao, W.C.H., Kulkarni, K., Zhang, Z., Kong, E.H., and Barford, D. (2012). Structure of the mitotic checkpoint complex. Nature 484, 208–213.

De Antoni, A., Pearson, C.G., Cimini, D., Canman, J.C., Sala, V., Nezi, L., Mapelli, M., Sironi, L., Faretta, M., Salmon, E.D., et al. (2005). The Mad1/Mad2 complex as a template for Mad2 activation in the spindle assembly checkpoint. Curr. Biol. CB 15, 214–225.

DeAntoni, A., Sala, V., and Musacchio, A. (2005). Explaining the oligomerization properties of the spindle assembly checkpoint protein Mad2. Philos. Trans. R. Soc. Lond. B. Biol. Sci. 360, 637–647, discussion 447-448.

Dobles, M., Liberal, V., Scott, M.L., Benezra, R., and Sorger, P.K. (2000). Chromosome missegregation and apoptosis in mice lacking the mitotic checkpoint protein Mad2. Cell 101, 635–645.

Eytan, E., Wang, K., Miniowitz-Shemtov, S., Sitry-Shevah, D., Kaisari, S., Yen, T.J., Liu, S.-T., and Hershko, A. (2014). Disassembly of mitotic checkpoint complexes by the joint action of the AAA-ATPase TRIP13 and p31comet. Proc. Natl. Acad. Sci. 111, 12019–12024.

Fang, G., Yu, H., and Kirschner, M.W. (1998). The checkpoint protein MAD2 and the mitotic regulator CDC20 form a ternary complex with the anaphase-promoting complex to control anaphase initiation. Genes Dev. 12, 1871–1883.

Habu, T., Kim, S.H., Weinstein, J., and Matsumoto, T. (2002). Identification of a MAD2-binding protein, CMT2, and its role in mitosis. EMBO J. 21, 6419–6428.

Hagan, R.S., Manak, M.S., Buch, H.K., Meier, M.G., Meraldi, P., Shah, J.V., and Sorger, P.K. (2011). p31(comet) acts to ensure timely spindle checkpoint silencing subsequent to kinetochore attachment. Mol. Biol. Cell 22, 4236–4246.

Hall, F.L., and Vulliet, P.R. (1991). Proline-directed protein phosphorylation and cell cycle regulation. Curr. Opin. Cell Biol. 3, 176–184.

Hanks, S., Coleman, K., Reid, S., Plaja, A., Firth, H., Fitzpatrick, D., Kidd, A., Méhes, K., Nash, R., Robin, N., et al. (2004). Constitutional aneuploidy and cancer predisposition caused by biallelic mutations in BUB1B. Nat. Genet. 36, 1159–1161.

Hara, M., Özkan, E., Sun, H., Yu, H., and Luo, X. (2015). Structure of an intermediate conformer of the spindle checkpoint protein Mad2. Proc. Natl. Acad. Sci. U. S. A. 112, 11252–11257.

Heo, W.D., Inoue, T., Park, W.S., Kim, M.L., Park, B.O., Wandless, T.J., and Meyer, T. (2006). PI(3,4,5)P3 and PI(4,5)P2 lipids target proteins with polybasic clusters to the plasma membrane. Science 314, 1458–1461.

Hingorani, S.R., Wang, L., Multani, A.S., Combs, C., Deramaudt, T.B., Hruban, R.H., Rustgi, A.K., Chang, S., and Tuveson, D.A. (2005). Trp53R172H and KrasG12D cooperate to promote chromosomal instability and widely metastatic pancreatic ductal adenocarcinoma in mice. Cancer Cell 7, 469–483.

Hochegger, H., Takeda, S., and Hunt, T. (2008). Cyclin-dependent kinases and cell-cycle transitions: does one fit all? Nat. Rev. Mol. Cell Biol. 9, 910–916.

Iwanaga, Y., Chi, Y.-H., Miyazato, A., Sheleg, S., Haller, K., Peloponese, J.-M., Li, Y., Ward, J.M., Benezra, R., and Jeang, K.-T. (2007). Heterozygous deletion of mitotic arrest-deficient protein 1 (MAD1) increases the incidence of tumors in mice. Cancer Res. 67, 160–166.

Kamata, T., and Pritchard, C. (2011). Mechanisms of aneuploidy induction by RAS and RAF oncogenes. Am. J. Cancer Res. 1, 955–971.

Kim, S., Sun, H., Ball, H.L., Wassmann, K., Luo, X., and Yu, H. (2010). Phosphorylation of the spindle checkpoint protein Mad2 regulates its conformational transition. Proc. Natl. Acad. Sci. U. S. A. 107, 19772–19777.

Lee, C.H., Della, N.G., Chew, C.E., and Zack, D.J. (1996). Rin, a neuron-specific and calmodulin-binding small G-protein, and Rit define a novel subfamily of ras proteins. J. Neurosci. Off. J. Soc. Neurosci. 16, 6784–6794.

Lengauer, C., Kinzler, K.W., and Vogelstein, B. (1997). Genetic instability in colorectal cancers. Nature 386, 623–627.

Li, M., Fang, X., Wei, Z., York, J.P., and Zhang, P. (2009). Loss of spindle assembly checkpoint-mediated inhibition of Cdc20 promotes tumorigenesis in mice. J. Cell Biol. 185, 983–994.

Lindqvist, A., Rodríguez-Bravo, V., and Medema, R.H. (2009). The decision to enter mitosis: feedback and redundancy in the mitotic entry network. J. Cell Biol. 185, 193–202.

Luo, X., Fang, G., Coldiron, M., Lin, Y., Yu, H., Kirschner, M.W., and Wagner, G. (2000). Structure of the Mad2 spindle assembly checkpoint protein and its interaction with Cdc20. Nat. Struct. Biol. 7, 224–229.

Luo, X., Tang, Z., Rizo, J., and Yu, H. (2002). The Mad2 spindle checkpoint protein undergoes similar major conformational changes upon binding to either Mad1 or Cdc20. Mol. Cell 9, 59–71.

Mapelli, M., Filipp, F.V., Rancati, G., Massimiliano, L., Nezi, L., Stier, G., Hagan, R.S., Confalonieri, S., Piatti, S., Sattler, M., et al. (2006). Determinants of conformational dimerization of Mad2 and its inhibition by p31comet. EMBO J. 25, 1273–1284.

Mapelli, M., Massimiliano, L., Santaguida, S., and Musacchio, A. (2007). The Mad2 conformational dimer: structure and implications for the spindle assembly checkpoint. Cell 131, 730–743.

Michel, L.S., Liberal, V., Chatterjee, A., Kirchwegger, R., Pasche, B., Gerald, W., Dobles, M., Sorger, P.K., Murty, V.V., and Benezra, R. (2001). MAD2 haplo-insufficiency causes premature anaphase and chromosome instability in mammalian cells. Nature 409, 355–359.

Mondal, G., Baral, R.N., and Roychoudhury, S. (2006). A new Mad2-interacting domain of Cdc20 is critical for the function of Mad2-Cdc20 complex in the spindle assembly checkpoint. Biochem. J. 396, 243–253.

Musacchio, A. (2015). The Molecular Biology of Spindle Assembly Checkpoint Signaling Dynamics. Curr. Biol. CB 25, R1002–1018.

Musacchio, A., and Salmon, E.D. (2007). The spindle-assembly checkpoint in space and time. Nat. Rev. Mol. Cell Biol. 8, 379–393.

Oromendia, A.B., and Amon, A. (2014). Aneuploidy: implications for protein homeostasis and disease. Dis. Model. Mech. 7, 15–20.

Pemble, H., Kumar, P., van Haren, J., and Wittmann, T. (2017). GSK3-mediated CLASP2 phosphorylation modulates kinetochore dynamics. J. Cell Sci. 130, 1404–1412.

Pino, L.K., Searle, B.C., Bollinger, J.G., Nunn, B., MacLean, B., and MacCoss, M.J. (2020). The Skyline ecosystem: Informatics for quantitative mass spectrometry proteomics. Mass Spectrom. Rev. 39, 229–244.

Primorac, I., and Musacchio, A. (2013). Panta rhei: the APC/C at steady state. J. Cell Biol. 201, 177–189.

Ran, F.A., Hsu, P.D., Wright, J., Agarwala, V., Scott, D.A., and Zhang, F. (2013). Genome engineering using the CRISPR-Cas9 system. Nat. Protoc. 8, 2281–2308.

Rodriguez-Viciana, P., Sabatier, C., and McCormick, F. (2004). Signaling specificity by Ras family GTPases is determined by the full spectrum of effectors they regulate. Mol. Cell. Biol. 24, 4943–4954.

Santaguida, S., Tighe, A., D’Alise, A.M., Taylor, S.S., and Musacchio, A. (2010). Dissecting the role of MPS1 in chromosome biorientation and the spindle checkpoint through the small molecule inhibitor reversine. J. Cell Biol. 190, 73–87.

Schilling, B., Rardin, M.J., MacLean, B.X., Zawadzka, A.M., Frewen, B.E., Cusack, M.P., Sorensen, D.J., Bereman, M.S., Jing, E., Wu, C.C., et al. (2012). Platform-independent and label-free quantitation of proteomic data using MS1 extracted ion chromatograms in skyline: application to protein acetylation and phosphorylation. Mol. Cell. Proteomics MCP 11, 202–214.

Schindelin, J., Arganda-Carreras, I., Frise, E., Kaynig, V., Longair, M., Pietzsch, T., Preibisch, S., Rueden, C., Saalfeld, S., Schmid, B., et al. (2012). Fiji: an open-source platform for biological-image analysis. Nat. Methods 9, 676–682.

Shao, H., and Andres, D.A. (2000). A novel RalGEF-like protein, RGL3, as a candidate effector for rit and Ras. J. Biol. Chem. 275, 26914–26924.

Simanshu, D.K., Nissley, D.V., and McCormick, F. (2017). RAS Proteins and Their Regulators in Human Disease. Cell 70, 17–33.

Sironi, L., Melixetian, M., Faretta, M., Prosperini, E., Helin, K., and Musacchio, A. (2001). Mad2 binding to Mad1 and Cdc20, rather than oligomerization, is required for the spindle checkpoint. EMBO J. 20, 6371–6382.

Sironi, L., Mapelli, M., Knapp, S., De Antoni, A., Jeang, K.-T., and Musacchio, A. (2002). Crystal structure of the tetrameric Mad1-Mad2 core complex: implications of a “safety belt” binding mechanism for the spindle checkpoint. EMBO J. 21, 2496–2506.

UniProt Consortium (2019). UniProt: a worldwide hub of protein knowledge. Nucleic Acids Res. 47, D506–D515.

Van, R., Cuevas-Navarro, A., Castel, P., and McCormick, F. (2020). The molecular functions of RIT1 and its contribution to human disease. Biochem. J. 477, 2755–2770.

Wall, V.E., Garvey, L.A., Mehalko, J.L., Procter, L.V., and Esposito, D. (2014). Combinatorial assembly of clone libraries using site-specific recombination. Methods Mol. Biol. Clifton NJ 1116, 193–208.

Wang, K., Sturt-Gillespie, B., Hittle, J.C., Macdonald, D., Chan, G.K., Yen, T.J., and Liu, S.-T. (2014). Thyroid Hormone Receptor Interacting Protein 13 (TRIP13) AAA-ATPase Is a Novel Mitotic Checkpoint-silencing Protein. J. Biol. Chem. 289, 23928–23937.

Wes, P.D., Yu, M., and Montell, C. (1996). RIC, a calmodulin-binding Ras-like GTPase. EMBO J. 15, 5839–5848.

Westhorpe, F.G., Tighe, A., Lara-Gonzalez, P., and Taylor, S.S. (2011). p31comet-mediated extraction of Mad2 from the MCC promotes efficient mitotic exit. J. Cell Sci. 124, 3905–3916.

Xia, G., Luo, X., Habu, T., Rizo, J., Matsumoto, T., and Yu, H. (2004). Conformation-specific binding of p31(comet) antagonizes the function of Mad2 in the spindle checkpoint. EMBO J. 23, 3133–3143.

Yang, G., Mercado-Uribe, I., Multani, A.S., Sen, S., Shih, I.-M., Wong, K.-K., Gershenson, D.M., and Liu, J. (2013). RAS promotes tumorigenesis through genomic instability induced by imbalanced expression of Aurora-A and BRCA2 in midbody during cytokinesis. Int. J. Cancer 133, 275–285.

Yang, M., Li, B., Tomchick, D.R., Machius, M., Rizo, J., Yu, H., and Luo, X. (2007). p31comet blocks Mad2 activation through structural mimicry. Cell 131, 744–755.

Yaoita, M., Niihori, T., Mizuno, S., Okamoto, N., Hayashi, S., Watanabe, A., Yokozawa, M., Suzumura, H., Nakahara, A., Nakano, Y., et al. (2016). Spectrum of mutations and genotype-phenotype analysis in Noonan syndrome patients with RIT1 mutations. Hum. Genet. 135, 209–222.

